# Modeling the Human Segmentation Clock with Pluripotent Stem Cells

**DOI:** 10.1101/562447

**Authors:** Mitsuhiro Matsuda, Yoshihiro Yamanaka, Maya Uemura, Mitsujiro Osawa, Megumu K. Saito, Ayako Nagahashi, Megumi Nishio, Long Guo, Shiro Ikegawa, Satoko Sakurai, Shunsuke Kihara, Michiko Nakamura, Tomoko Matsumoto, Hiroyuki Yoshitomi, Makoto Ikeya, Takuya Yamamoto, Knut Woltjen, Miki Ebisuya, Junya Toguchida, Cantas Alev

## Abstract

Pluripotent stem cells (PSCs) have increasingly been used to model different aspects of embryogenesis and organ formation^1^. Despite recent advances in the *in vitro* induction of major mesodermal lineages and mesoderm-derived cell types^2,3^, experimental model systems that can recapitulate more complex biological features of human mesoderm development and patterning are largely missing. Here, we utilized induced pluripotent stem cells (iPSCs) for the stepwise *in vitro* induction of presomitic mesoderm (PSM) and its derivatives to model distinct aspects of human somitogenesis. We focused initially on modeling the human segmentation clock, a major biological concept believed to underlie the rhythmic and controlled emergence of somites, which give rise to the segmental pattern of the vertebrate axial skeleton. We succeeded to observe oscillatory expression of core segmentation clock genes, including *HES7* and *DKK1*, and identified novel oscillatory genes in human iPSC-derived PSM. We furthermore determined the period of the human segmentation clock to be around five hours and showed the presence of dynamic traveling wave-like gene expression within *in vitro* induced human PSM. Utilizing CRISPR/Cas9-based genome editing technology, we then targeted genes, for which mutations in patients with abnormal axial skeletal development such as *spondylocostal dysostosis* (SCD) (*HES7*, *LFNG* and *DLL3*) or *spondylothoracic dysostosis* (STD) (*MESP2*) have been reported. Subsequent analysis of patient-like iPSC knock-out lines as well as patient-derived iPSCs together with their genetically corrected isogenic controls revealed gene-specific alterations in oscillation, synchronization or differentiation properties, validating the overall utility of our model system, to recapitulate not only key features of human somitogenesis but also to provide novel insights into diseases associated with the formation and patterning of the human axial skeleton.

---

We initially aimed to mimic and recreate *in vitro* the signaling events responsible for the step-wise emergence of PSM and its derivatives during embryonic development, as also recently attempted by others^2,4,5^, via selective activation or inhibition of appropriate signaling pathways, using human iPSCs as starting material (Fig. 1a). We characterized the ability of our *in vitro* induced human PSM cells to differentiate into somitic mesoderm and its two main derivatives, sclerotome and dermomyotome, which give rise to bone and cartilage of the axial skeleton and skeletal muscle and dermis of the emerging embryo respectively. RNA-sequencing (RNA-seq) analysis and subsequent characterization of *in vitro* derived human PSM samples revealed, that at each step of our induction and differentiation protocol, markers expected to be present based on either embryological studies in animal models or recent reports utilizing stem cells^2,4–6^, were appropriately expressed at both transcript and protein levels (Fig. 1b-f, Extended Data Fig. 1a-f, Extended Data Table 1), indicating that our step-wise approach is following the developmental trajectory and recapitulating ontogeny seen during embryonic somitic mesoderm development.

**Fig. 1.**
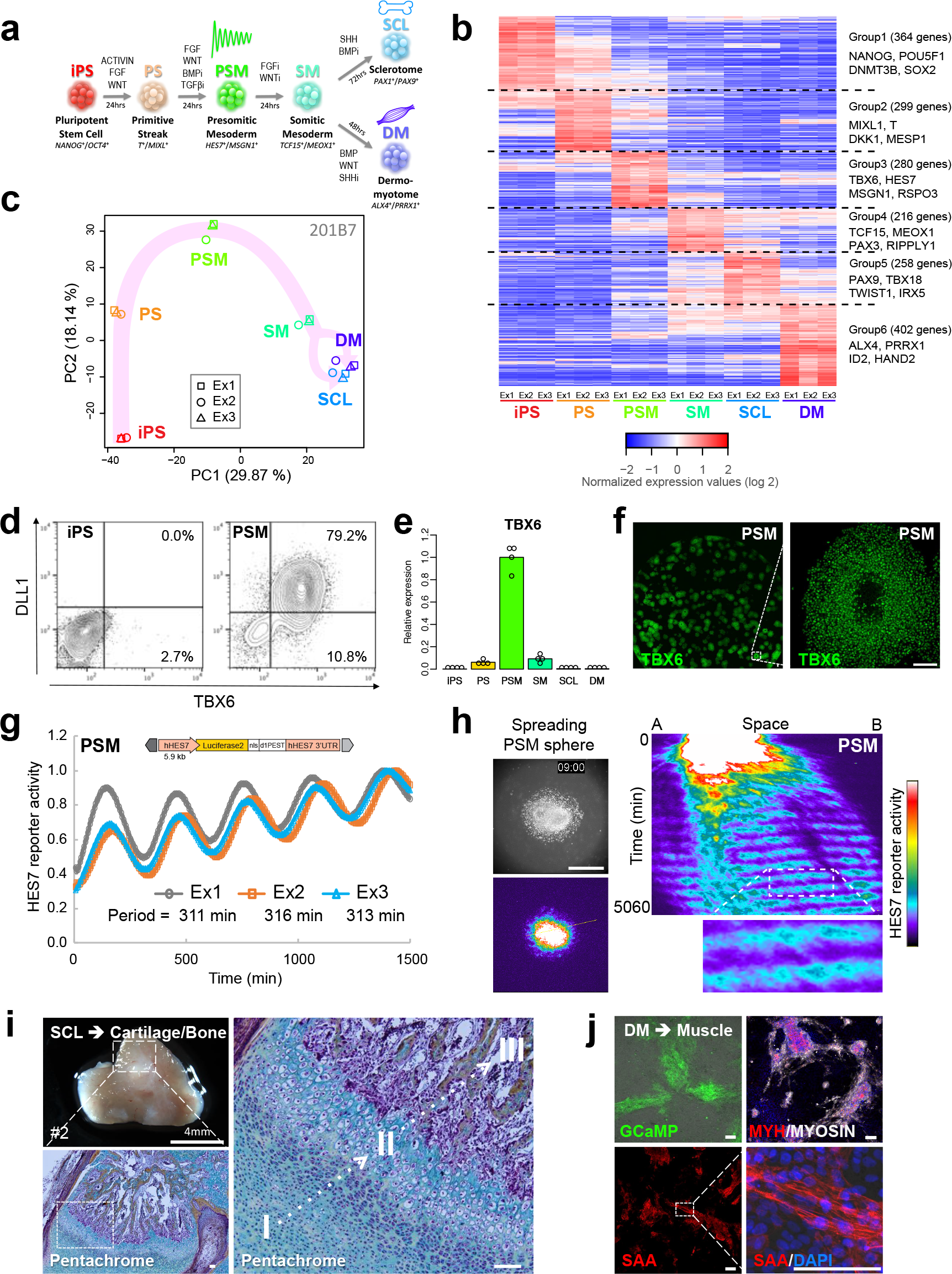
Molecular and functional analysis of human PSC-derived presomitic mesoderm (PSM) and its derivatives. **a**, Schematic overview of step-wise induction and differentiation of human PSM mimicking embryonic development via activation or inhibition of appropriate signaling pathways. **b**, Heatmap of gene expression levels of step-wise induced PSM and derivatives. FPKM values of each gene were normalized to mean of all samples. Results shown for samples derived from iPSC-line 201B7. See Extended Data Fig. 1 for data obtained for other iPSC-line (1231A3) and Extended Data Table 1 for complete list of identified genes. **c**, PCA analysis of transcript levels at different stages of applied protocol (for 201B7-derived samples). **d**, Protein expression of DLL1 and TBX6 in human iPSCs and iPSC-derived PSM (201B7). **e**, Expression of *TBX6* transcript during different stages of human PSM induction and differentiation (qRT-PCR results for four independent experiments using 201B7 are shown). **f**, Expression of TBX6 protein in human PSM. Left panel shows TBX6 staining of entire well. Enlarged view of highlighted area is shown on right side of panel. Scale bar: 100 µm. **g**, Oscillation of HES7-reporter activity in induced PSM. The signal was normalized to the maximum oscillation peak. The period was calculated as the average elapsed time between the peaks. Data of three independent experiments are shown. Schematic depiction of utilized reporter construct shown on top. **h**, Synchronization of HES7-reporter activity in spreading PSM sphere. Left: A PSM sphere was induced in 3D culture, before attaching to dish. Right: A kymograph along the yellow line in the left panel. Normalization (1500, 9000). Scale bar: 500 µm. Representative images of three independent experiments are shown. See also Extended Data Movie 1. **i**, Representative whole mount (upper panel) and histological staining of section (lower panel) of human sclerotome (SCL)-derived *in vivo* cartilage and bone. Pentachrome staining of marked area reminiscent of *in vivo* human endochondral bone formation. I: proliferative human cartilage, II: hypertrophic cartilage, III: ossifying cartilage and forming human bone. Scale bar: 100 µm. See also Extended Data Fig. 2e. **j**, Skeletal muscle derivation from step-wise induced human dermomyotome (DM). Left upper side of panel: GCaMP positive beating colonies of induced skeletal muscle. Left lower and right side of panel: Staining of *in vitro* induced human skeletal muscle with myosin heavy chain (MYH) and myosin (upper panel) or sarcomeric alpha-actinin (SAA) (lower half of panel). Scale bar: 100 µm. See also Extended Data Fig. 2f and Extended Data Movie 2.

We detected in our *in vitro* derived human PSM samples expression of *TBX6* and *DLL1*, two well-established markers of presomitic mesoderm^7^, at transcript and protein levels (Fig. 1d-f, Extended Data Fig. 1d-f). We also detected high-level expression of *HES7*, a known regulator of the segmentation clock in murine PSM^8^ (Fig. 1b, Extended Data Fig. 1c). Based on this observation we generated a luciferase-based iPSC reporter line for human *HES7* promoter activity (HES7-reporter). We observed clear oscillation of the HES7-reporter in our human PSM samples (Fig. 1g) and determined the period of the human segmentation clock to be 5-6 hours, which is similar to the 4-6 hour period reported for somite formation in primary human embryo samples^9,10^. To our knowledge, this is the first real-time observation of oscillatory expression in the human segmentation clock.

We then asked, whether we could also observe traveling wave-like expression, another hallmark of the segmentation clock, which is caused by the synchronization among oscillations in neighboring cells. Such traveling waves have been reported in the context of explant studies utilizing reporter mice and mouse ESC-derived PSM^11,12^, but have never been observed in human PSM. Using a sphere of human PSM induced in 3D culture, we could see sustained oscillation and the clear presence of traveling waves (Extended Data Movie 1), which is also indicated by the tilted slope in the kymograph (Fig. 1h).

In order to ensure that our *in vitro*-derived PSM is indeed comparable to its *in vivo* counterpart, we further characterized its differentiation capacity into somitic mesoderm, sclerotome and dermomyotome. To induce somitic mesoderm we mimicked the decrease in Fgf and Wnt activity along the posterior-anterior axis of the PSM reported in the embryonic context^13^, by simultaneous inhibition of both pathways, leading to the rapid and robust induction of somitic mesoderm expressing *TCF15*, a well established marker of somite development^14^, at the transcript (Fig. 1b, Extended Data Fig. 1c) and the protein level (Extended Data Fig. 1g). In addition, *MESP2*, a marker of segmentation during somitogenesis, showed localized albeit weak expression in induced human somitic mesoderm, similar to the remnant expression of *TBX6* at this stage, indicating that, despite a strong signal for differentiation, patterning reminiscent of somite segmentation might take place *in vitro*, as recently shown for mouse ESC-derived PSM^12^ (Extended Data Fig. 1g, right side of panel). Furthermore, dermomytome and sclerotome cells derived from *in vitro* induced human somitic mesoderm expressed appropriate developmental stage-specific markers at the transcript and protein level (Extended Data Fig. 1c, 2a). To further demonstrate that our induced human somitic mesoderm and the sclerotome derived thereof, is, indeed functional and can give rise to cartilage and bone, as seen during embryogenesis^15^, we performed *in vitro* and *in vivo* differentiation and transplantation assays (Extended Data Fig. 2b-e). As also recently reported by others^4^, we could see self-organization of bone and cartilage, including properly patterned endochondral bone, from transplanted human sclerotome cells *in vivo* (Fig. 1i, Extended Data Fig. 2b, 2c, 2e). We then examined our *in vitro* induced dermomyotome cells for their ability to give rise to human skeletal muscle *in vitro*. We could observe robust induction of skeletal muscle cells, showing expression of proper muscle markers at the protein level (Fig. 1j, Extended Data Fig. 2f). Functional analysis of dermomyotome cells derived from a calcium reporter iPSC line (Gen1C) revealed the reproducible presence of beating skeletal muscle cells after three weeks of *in vitro* 2D differentiation culture (Fig. 1j, Extended Data Movie 2). We thus showed that our *in vitro* step-wise induction protocol can give rise to human PSM cells and their proper derivatives that appear to recapitulate key features of their corresponding developmental counterparts and respective embryological stages.

Since the oscillation and synchronization in human PSM have never been characterized before, we asked whether we could deepen our understanding of the human segmentation clock even further, using the experimental system at hand. To this end we collected PSM samples during oscillation by monitoring the oscillatory activity of the HES7*-*reporter (Extended Data Fig. 3a) and performed RNA-seq analysis of the isolated RNA samples (16 samples each for two independent sets of experiments).

Our NGS-based analysis of the different PSM time-points revealed a core-set of about two hundred oscillating genes (Fig. 2a, Extended Data Table 2). Pathway and gene ontology (GO) analysis of the identified gene clusters revealed that in addition to enrichment of pathway members previously associated with the segmentation clock, such as Notch, Wnt or Fgf signaling^16,17^ novel pathways were also represented in our data set, including members of the TGFβ pathway or Hippo signaling (Fig. 2b, Extended Data Fig. 3b, Extended Data Table 3). Interestingly, the Hippo pathway component YAP was recently reported to be an important regulator of oscillatory activity in mouse PSM^18^, suggesting that this role might be conserved in human PSM, which showed oscillatory activity of core Hippo pathway components (*TEAD4*, *AMOTL2*) (Fig. 2a, 2b, Extended Data Fig. 3b). We also detected several HOX genes showing oscillatory expression in human PSM (*HOXA3, HOXA5*, *HOXD1*) (Fig. 2a, 2b, Extended Data Fig. 3b). Oscillatory expression of *HOXD1* was previously shown during somitogenesis in mouse^19^ suggesting that its expression pattern and biological function might be conserved in human PSM. We could also identify oscillation of putative modulators of the cytoskeleton (*ARHGAP24*, *ARHGEF2*, *RHOU*, *PLEKHG2*) as well as histone modifiers (*KDM6B*, *JADE1*) as oscillating genes (Fig. 2a, 2c, Extended Data Fig. 3b). The intriguing possibility that above-mentioned cytoskeleton associated oscillating genes might represent a link between the segmentation clock and the actual process of segmentation, characterized by mesenchymal to epithelial transition and associated with major cytoskeletal rearrangements, remains to be elucidated. Furthermore, identification of oscillating histone modifiers in the PSM, suggests a possible role of epigenetic modifications in the regulation or maintenance of the segmentation clock, and will be the topic of future research efforts.

**Fig. 2.**
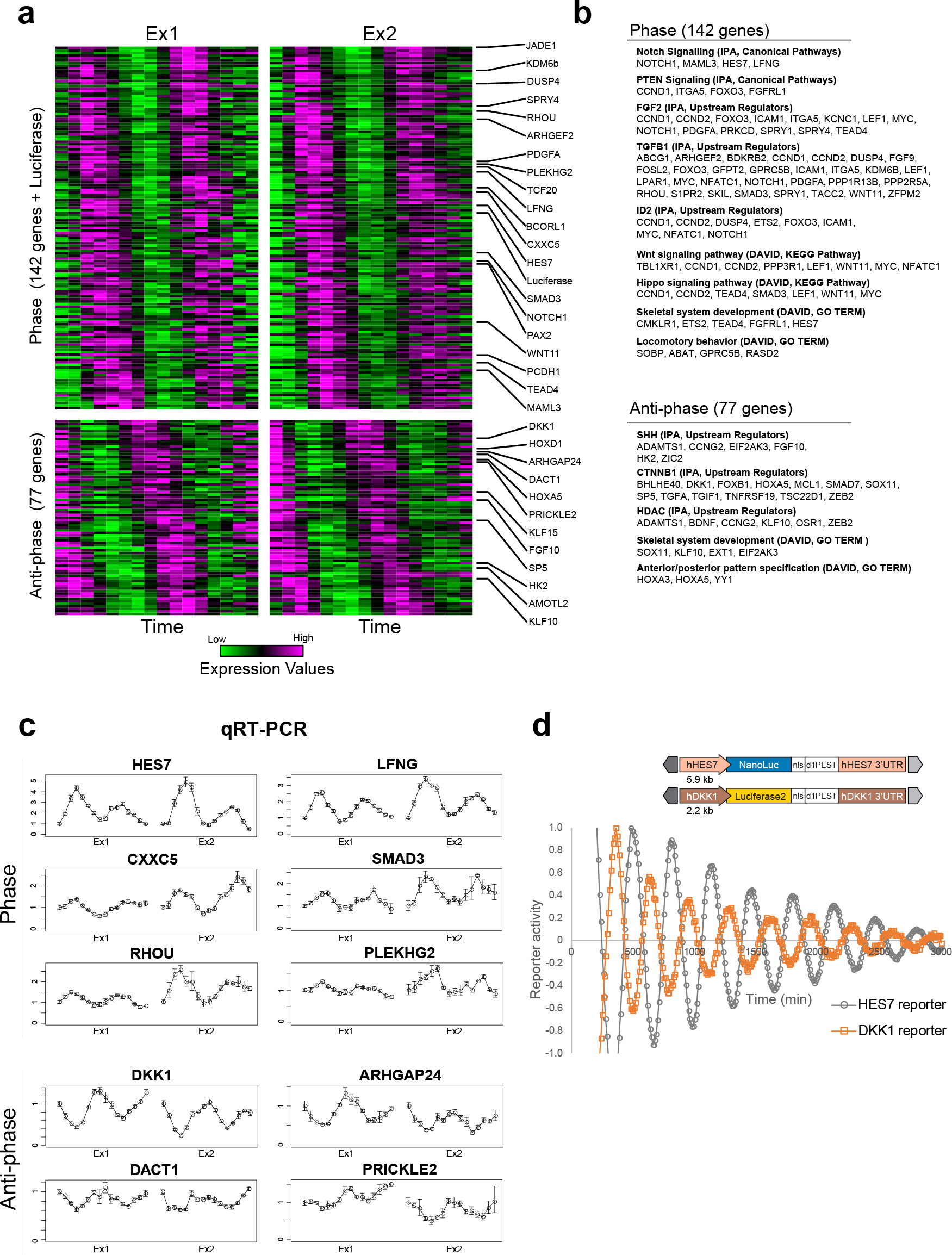
Identification of phase and anti-phase oscillating genes of the human segmentation clock. **a**, Heatmap of normalized gene expression levels for oscillating genes in human PSM. RNA-sequencing results for two independent data sets, with 16 samples each. Examples of identified phase and anti-phase oscillating genes are highlighted on right side of panel. See also Extended Data Table 2 for complete list of identified oscillating genes. **b**, Pathway analysis result of phase and anti-phase oscillating genes. See also Extended Data Table 3 for complete results of pathway and GO analyses. **c**, Validation of RNA-sequencing results via qRT-PCR for selected phase and anti-phase oscillating genes. Error bars indicate S.D. of three technical replicas for each time point and sample set. **d**, Results obtained for dual luciferase reporter assay of HES7-reporter (NanoLuc) and DKK1-reporter (Luciferase2) shown in lower half of panel. The one-step induction protocol was used. The signal was detrended (± 2 hours) and normalized to the maximum oscillation peak. Representative graph of three independent experiments is shown. Schematic overview of used reporter constructs are shown on top.

Among the identified oscillatory genes, two thirds were oscillating in phase with *HES7* (142) and about one third (77) showing an anti-phase oscillatory expression pattern (Fig. 2a, Extended Data Table 2). The phase cluster of human oscillating genes contained Notch-pathway associated genes such as *LFNG*^20^, while the anti-phase cluster contained Wnt-pathway associated negative feedback regulators such as *DKK1* and *SP5*^21,22^ (Fig. 2a, 2c), as previously reported for posterior PSM of mouse embryos^23^. Generating a dual luciferase-activity based reporter line for *DKK1* and *HES7* promoter activities we confirmed clear phase and anti-phase reporter oscillations in human iPSC-derived PSM samples (Fig. 2d), suggesting that our induced PSM may represent posterior immature PSM rather than anterior mature PSM.

In order to also show the utility of our system to model anomalies during human axial skeletogenesis such as SCD or STD, known to be caused by mutations in genes associated with the segmentation clock (e.g. *HES7*, *LFNG*, *DLL3* and *MESP2*)^24^, we utilized CRISPR/Cas9-mediated genome editing technology to generate knock-out reporter iPSC lines aiming to induce frameshifts or deletion mutations in these target genes in the *HES7* luciferase reporter line background (Extended Data Fig. 4) and analyzed their putative loss-of-function effect on oscillatory HES7-reporter activity. Knock-out of endogenous *HES7* itself led to complete loss of oscillatory activity of the HES7-reporter, in a 2D-oscillation assay (Fig. 3a), similar to previous embryological studies utilizing knock-out mice^25^. Interestingly knock-out reporter lines for *LFNG*, *DLL3* or *MESP2* continued to show strong oscillatory *HES7* activity (Fig. 3b) even though knock-out mice for *LFNG* and *DLL3* were reported to show defective oscillation patterns^20,26^. We reasoned that in this 2D-oscillation assay the phase (i.e. timing) of oscillations is initially reset by medium change, showing collective oscillation even in the absence of a strong synchronization mechanism. We then examined the synchronization ability of knock-out reporter lines of aforementioned genes using 3D-spheres of PSM (Fig. 3c). The control (healthy/wild-type) and the knock-out reporter line for *MESP2* showed sustained oscillations and occasional traveling waves (Extended Data Movie 3), indicating intact synchronization among neighboring cells. In the knock-out lines of *LFNG* or *DLL3*, by contrast, oscillation damped quickly and clear traveling waves were not observed (Extended Data Movie 3, Fig. 3c, 3d, Extended Data Fig. 5a). We interpreted the observed quick loss of oscillatory activity as a sign of diminished synchronization. Unlike the 2D-oscillation assay, spheres spread dynamically on the culture dish in the 3D-synchronization assay, and cell movement desynchronizes the oscillation phases. Without proper synchronization mechanism collective oscillation is quickly lost even though oscillations in individual cells continue. Thus, our assay systems using induced human PSM can detect defects in both oscillation and synchronization.

**Fig. 3.**
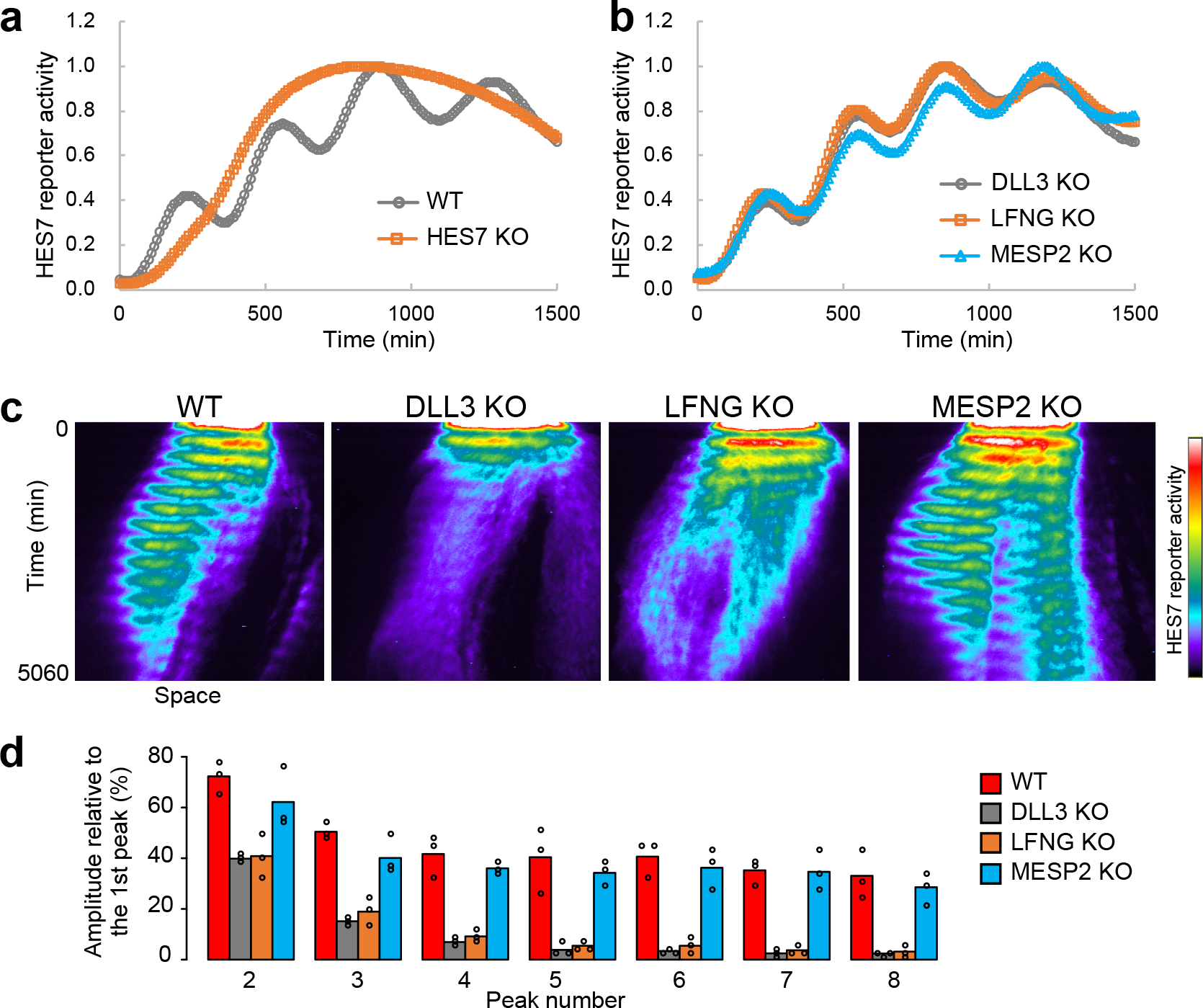
Molecular and functional evaluation of targeted disruption of selected segmentation clock genes in human PSM. **a**, 2D-oscillation assay for wild-type (WT) and *HES7*-knock-out PSMs. The signal was normalized to the maximum oscillation peak. **b**, 2D-oscillation assay for *LFNG*-, *DLL3*- and *MESP2*-knock-out PSMs. **c**, 3D-synchronization assay for knock-out PSMs. A kymograph along the yellow line in Extended Data Movie 3 is shown. Normalization (1500, 35000). Representative graphs and images of three independent experiments are shown. **d**, Damping rate of oscillation amplitude in knock-out PSMs. The signal value of each oscillation peak is shown. Data of three independent experiments are shown. See also Extended Data Fig. 5a.

Flow cytometric evaluation as well as RNA-seq based transcriptome analysis showed no major differences between control cells and aforementioned reporter lines (Extended Data Fig. 5b, 5c). PSM induction efficiency was high and comparable to wild-type (healthy donor) cells in *DLL3*, *LFNG* and *MESP2* knock-out reporter lines, while slightly reduced for *HES7* knock-out iPSC lines (Extended Data Fig. 5b). Few differences in gene expression at the iPSC and PSM stages were observed when comparing knock-out lines with the original healthy donor line, with *HES7, MESP2* and *LFNG* showing higher expression in *HES7* knock-out derived PSM, as previously also shown in mice^27^ (Extended Data Fig. 5c).

Taken together, these results underline the overall value of an higher order assay system that can assess not only gene or protein expression but also more complex features such as oscillation or synchronization in human PSM, thus opening up the path to work out functionally relevant and possibly disease-associated features specific to each loss- or gain-of-function mutation, otherwise not accessible.

To further evaluate the utility of our experimental model system to assess not only key features of the human segmentation clock and somitogenesis but also address causative molecular mechanisms associated with diseases affecting axial skeletogenesis, we established iPSC lines from 12 individuals afflicted by SCD or STD (data not shown). In one of the established STD patient-derived iPSC lines (Extended Data Fig. 6) we identified via Exome sequencing compound heterozygous loss-of-function mutations in *MESP2* (rs1452984345: c.256delGCCA, p.fsTer118; rs71647808: c.307T, p.E103X) (Fig. 4a). Induction of PSM from these patient iPSCs (STD-A and STD-F; two different iPSC clones from the same patient) appeared to be not affected, as assessed by flow cytometric analysis of DLL1 expression (Fig. 4b; Extended Data Fig. 7a). We observed for the patient line (STD-A) harboring *MESP2* loss-of-function mutations, clear oscillatory activity of *HES7* in 2D-oscillation assay (Fig. 4c), similar to what we saw for our human *MESP2* knock-out reporter lines (Fig. 3b). In the 3D-synchronization assay this patient line also showed sustained collective oscillation and occasional traveling waves, indicating an intact synchronization mechanism (Fig. 4d), again similar to results seen for *MESP2* knock-out lines (Fig. 3c). In order to facilitate the molecular and functional analysis of our patient lines, we generated isogenic controls by correcting the underlying putative disease causing mutations via gene targeting with CRISPR/Cas9. Allele-specific gene correction of *MESP2* was achieved using sgRNAs targeting either the c.256delGCCA or c.307T mutation and homologous recombination with donor vectors bearing normal *MESP2* gene sequence. Microhomology-assisted excision (MhAX) was used to remove the selection cassette^28^ (Fig. 4e, 4f, Extended Data Fig. 8), thus effectively rescuing the disease-causing loss of *MESP2*, albeit heterozygously. Gene-edited iPSCs were confirmed to be karyotypically similar to the parental patient iPSC line (Extended Data Fig. 9). As no clear oscillation or synchronization phenotype could be observed for the analyzed patient line, we asked whether we could see possible differences at the functional or molecular level by comparing patient (STD-A and STD-F) and corresponding rescued iPSC lines (STD-resA and STD-resF). To this end we induced and compared all stages of our *in vitro* induction and differentiation protocol via RNA-seq analysis using patient and rescue lines (Fig. 4g, Extended Data Fig. 7b).

**Fig. 4.**
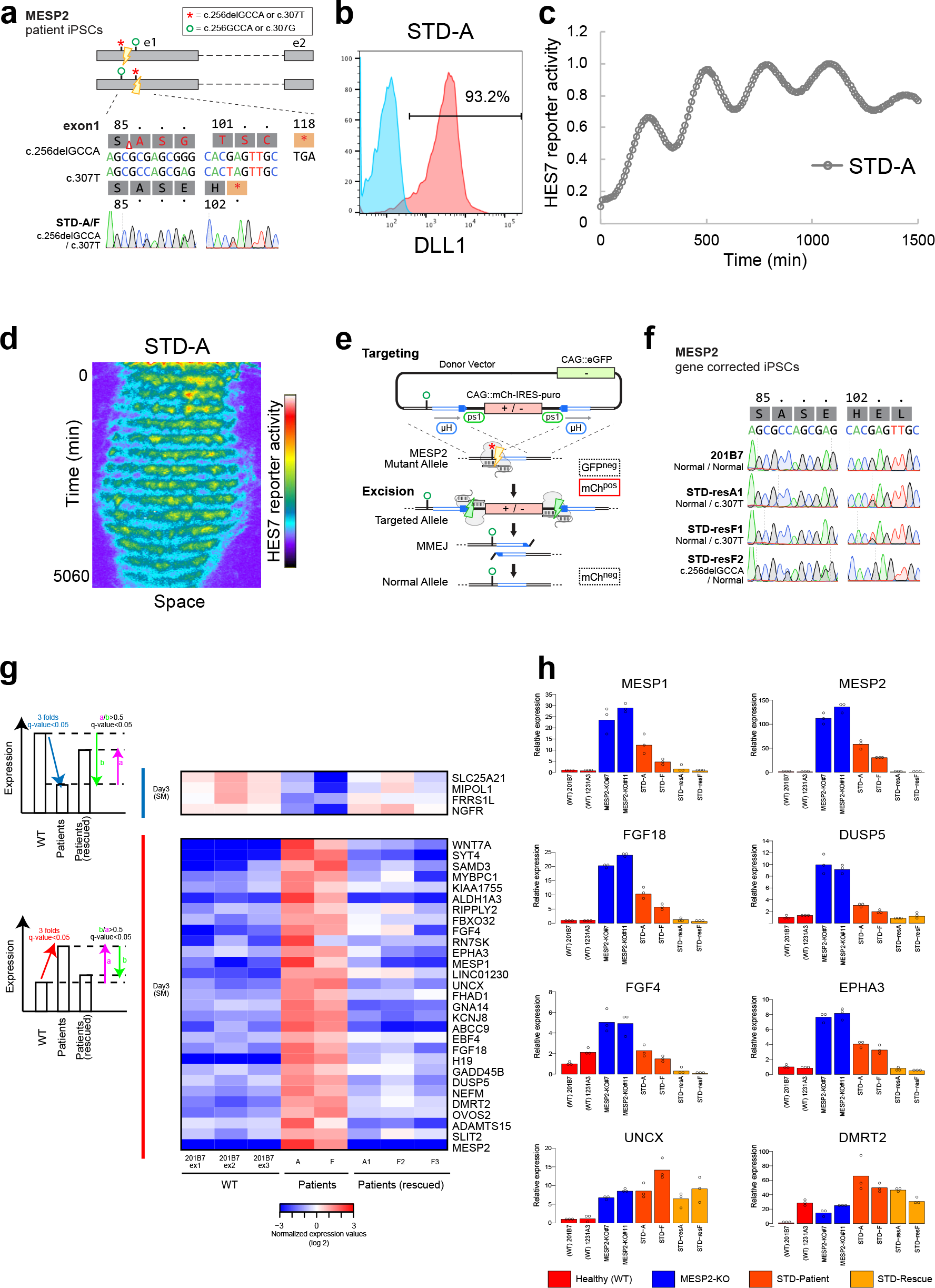
*In vitro* recapitulation and molecular analysis of disease-phenotype using spondylothoracic dysostosis (STD) patient iPSCs and genetically corrected controls. **a**, Compound heterozygous genotype of STD-A/F patient iPSCs. **b**, Evaluation of DLL1 positive PSM induction efficiency of STD-A patient iPSCs. Representative results of DLL1 protein expression in STD-A derived PSM, n=3. See also Extended Data Fig. 7a for representative results of comparison (n=3) between patient (STD-A/F) and rescue clones (STD-resA/resF). **c**, 2D-oscillation assay for STD-A PSM. The signal was normalized to the maximum oscillation peak. Representative graph of three independent experiments is shown. **d**, 3D-synchronization assay for STD-A PSM. Normalization (1500, 4000). Representative image of three independent experiments is shown. **e**, Schematic of the gene targeting procedure for allele-specific correction of *MESP2* mutations using MhAX. Details for the targeting and genotyping procedures are provided in Extended Data Fig. 8. **f**, Genotype of heterozygously corrected iPSC subclones. 201B7 is included as a reference. **g**, Heatmap of gene expression levels for STD-related genes. The genes were upregulated or downregulated in the STD-A and STD-F patient lines and increases or decreases were inhibited in rescued cell lines. See also Extended Data Fig. 7b. **h**, qRT-PCR-based validation of RNA-sequencing results. Data of three independent experiments are shown. See also Extended Data Fig. 7c for qRT-PCR-based validation of additional candidates.

Comparison of patient clones with wild-type (healthy) and heterozygously corrected lines revealed the presence of an up-regulated gene cluster at the somitic mesoderm stage in the analyzed patient lines, which could be reversed upon correction of either mutated allele (Fig. 4g). Genes apparently up-regulated in patient somitic mesoderm and reduced upon rescue of either *MESP2* mutation, included *FGF4*, *FGF18* and *DUSP5* (Fig. 4g, 4h), indicating abnormal Fgf signaling as a possible novel disease associated molecular feature in STD. *MESP2* knock-out iPSC-derived somitic mesoderm samples also showed higher levels of expression of *FGF4*, *FGF18* and *DUSP5* (Fig. 4h). Interestingly human *EPHA3*, which was previously reported to have a dominant negative effect on somite formation and axial organization in fish^29^, was also found to be up-regulated in STD-patient and MESP2-knock-out derived somitic mesoderm (Fig. 4g, 4h). Knock-out and patient-lines showed further higher levels of expression for *MESP1* and *MESP2* as compared to the levels found in healthy or genetically corrected control samples, indicating possible disrupted negative feed-back regulation by human *MESP2*. *UNCX*, a gene associated with rostro-caudal patterning of forming somites and found to be up-regulated during abnormal somitogenesis in *MESP2* knock-out mice^30^ was also found to be highly expressed in *MESP2* patient iPSC-derived somitic mesoderm samples as compared to genetically rescued controls. Interestingly, several other genes associated with patterning during somitogenesis and for which genetic mutations in SCD patients were recently reported, including *LFNG*, *RIPPLY2* and *DMRT2^24^* were also up-regulated in somitic mesoderm samples derived from STD patient lines (STD-A and STD-F) harboring *MESP2* loss-of-function mutations (Fig. 4g, 4h, Extended Data Fig. 7c), indicating reciprocal regulatory mechanisms, possibly connecting these disease associated genes at the molecular and functional level during the pathogenesis of axial skeletogenic abnormalities including SCD and STD.

In summary, we have shown for the first time phase- and anti-phase oscillation and traveling wave-like gene expression of key segmentation clock genes in human PSM, and identified a putative molecular network of known and novel members comprising the mammalian segmentation clock. We furthermore assessed the function of several disease-associated members of the human segmentation clock, applying our experimental model system in combination with patient-like or patient-derived iPSCs, thus effectively creating the first human PSC-based model for congenital scoliosis, which may contribute to decipher the molecular mechanisms underlying normal and pathological human axial skeletogenesis.

Furthermore, being able to induce iPSCs from non-human model systems including evolutionary distant species, may, in combination with our proposed experimental model system, also pave the way for interspecies comparisons of segmentation clocks, and address as yet elusive molecular and evolutionary biological questions, such as the presence or absence of an evolutionary conserved core network of segmentation clock genes. Having access to a robust experimental model system that can be easily manipulated without the need for transgenic animals or primary tissues, while allowing assessment of not only genetic but also environmental or epigenetic factors, will undoubtedly facilitate our molecular and functional understanding of proper as well as abnormal human somitogenesis and axial skeletal development.

## ACKNOWLEDGEMENTS

The authors would like to thank Brendan McIntyre and Paul O’Neill for critical reading of the manuscript; Kanae Mitsunaga for help with FACS analysis; Yuhei Ashida for help with development of 3D-synchronization assay; Junya Asahira for help with RNA-seq experiments; Akihiro Yamashita for help with 3D-CI experiments; Mitsuaki Shibata and Taiki Nakajima for help with development of one-step PSM induction protocol; Noriaki Kawakami for help with general information on SCD; Monika Ohno and Sayaka Nishimura for help with iPSC quality control and validation; the CiRA Genome Evaluation Group, in particular Hiromi Dohi, Fumiyo Kitaoka, Masaki Nomura, Tomoko Takahashi, Masafumi Umekage, and Naoko Takasu for performing SNP array analysis. This work was supported by the CiRA Fellowship Program of Challenge to C.A.; Naito Foundation Scientific Research Grant to C.A.; Grant-in-Aid for Challenging Exploratory Research (KAKENHI Number 16K15664) to C.A.; Grant-in-Aid for Scientific Research on Innovative Areas (KAKENHI Number 17H05777) to M.M.; Takeda Science Foundation Grant to M.E.; Japan Agency for Medical Research and Development (AMED) Grants Number 12103610 and 17935423 to M.K.S. for iPSC generation and qualification; the Core Center for iPS Cell Research and the Acceleration Program for Intractable Disease Research Utilizing Disease Specific iPS Cells (AMED) to J.T.; the Cooperative Research Program (Joint Usage/Research Center Program) of Institute for Frontier Life and Medical Sciences, Kyoto University to J.T., L.G and S.I..

## AUTHOR CONTRIBUTIONS

C.A. conceived the study; C.A. and M.E. designed the project and supervised it; M.M. developed and performed 2D-oscillation assay and 3D-synchronization assay; Y.Y., M.U. and C.A. developed induction and differentiation protocols and performed majority of remaining *in vitro* and *in vivo* experiments; S.K. supported microscopy and calcium imaging; M.Ni. helped with xeno-transplantation experiments; M.O., M.K.S. and A.N. established patient iPSC lines used in this study and performed quality control of iPSCs; M.O. helped with FACS data analysis; L.G. and S.I. performed exome sequencing and database analysis; T.Y. analyzed RNA-seq and qRT-PCR-data; S.S. helped with RNA-seq analysis; K.W. designed gene knockout and editing strategies; T.M. performed gene editing and genotyping of patient iPSCs; M.Na. performed genotyping of patient and gene-edited iPSCs; Y.Y., M.U. and C.A. generated knock-out lines with the help of M.Na., K.W. and performed molecular/functional assays using knock-out lines, patient iPSCs and corrected controls; M.I. developed one-step PSM induction protocol; M.K.S. and H.Y. shared reagents; J.T. helped with establishment of patient lines and provided administrative support; C.A. analyzed and interpreted the data and wrote the paper with the support of M.E. and K.W..

## AUTHOR INFORMATION

The authors declare no competing interests. Correspondence and request for materials should be addressed to M.E. or C.A.

## METHODS

### Pluripotent stem cell generation and culture

Experiments were performed using mainly two human induced pluripotent stem (iPS) cell lines derived from healthy donors, 1231A3^31^ and 201B7^32^. Pluripotent stem cells of patients suffering from *spondylocostal dysostosis* (SCD) or *spondylothoracic dysostosis* (STD) were induced using patient-derived primary cell samples. The following cell line of a STD patient was obtained from the NIGMS Human Genetic Cell Repository at the Coriell Institute for Medical Research: GM13539. Reprogramming was performed with episomes (pCE-hOCT3/4, pCE-hSK, pCE-hUL, pCE-mp53DD, pCXB-EBNA1) under feeder-free conditions using StemFit^®^ medium and laminin-coated dishes (iMatrix511)^31^. Human iPSCs were maintained without feeder cells and cultured on iMatrix-511 silk (Nippi) coated dishes or plates with StemFit^®^ AK02N (Ajinomoto) medium supplemented with 50 U penicillin and 50 mg/ml streptomycin (Gibco).

### Step-wise induction of human somitic mesoderm

Human iPSCs were seeded on iMatrix-511 silk-coated plates or dishes at appropriate densities as single cells (e.g. 1.3 × 10^4^ cells/well into 6 well plate; 8.0 × 10^4^ cells/dish into 10 cm dish) 5 days before induction. All differentiation and induction steps were performed in chemically defined medium with insulin (CDMi)^33^ if not otherwise mentioned. Our step-wise protocol is similar to a recently published mesoderm induction protocol^4^, albeit with some differences. Human primitive streak (PS) cells were induced by treatment of iPSCs with bFGF (20 ng/ml), CHIR99021 (10 µM) and Activin A (50 ng/ml) for 24 hours. Presomitic mesoderm (PSM) cells were induced from PS cells by exposure to SB431542 (10 µM), CHIR99021 (3 µM), LDN193189 (250 nM) and bFGF (20 ng/ml) for 24 hours. Subsequently, somitic mesoderm (SM) cells were induced from PSM cells using PD173074 (100 nM) and XAV939 (1 µM) for 24 hours. For details of used recombinant human proteins and small molecule agonists or inhibitors see Extended Data Table 4.

### Human sclerotome and dermomyotome induction

Following initial step-wise somitic mesoderm (SM) induction, human sclerotome (SCL) cells were induced with combination of SAG (100 nM) and LDN193189 (600 nM)^34^ for 72 hours. Dermomyotome (DM) cells were induced from human somitic mesoderm as previously described^4^, using a combination of CHIR99021 (3 µM), GDC0449 (150 nM) and BMP4 (50 ng/ml) for 48 hours.

### *In vitro* 3D-chondrogenic induction (3D-CI)

Step-wise induced human sclerotome (SCL) cells were dissociated using Accutase (Life Tech), centrifuged and resuspended in CDMi before being seeded (2.0 × 10^5^ cells/well) into 96 well low attachment plates containing sclerotome induction medium with ROCK-inhibitor Y27632 (Wako), forming 3D aggregates overnight. Initial 3D-SCL spheres were transferred into low attachment plates or dishes containing 3D-CI medium^35^ and cultured under standard conditions. Medium was changed every three days.

### *In vitro* skeletal muscle induction

Dermomyotome (DM) cells were dissociated using Accutase (Life Tech), centrifuged, resuspended in CDMi and seeded (2.5 × 10^5^ cells/well) into Matrigel coated 12 well plates in muscle induction medium containing ROCK-inhibitor Y27632 (Wako). In order to induce human skeletal muscle cells, we applied the N2 medium established previously^36^ with some modifications (DMEM/F12 (Gibco), 1% ITS (Corning), 1% N2 Supplement (Gibco), 0.2% Penicillin/Streptomycin (Gibco), 1% L-Glutamine (Gibco), 2% Horse serum (Sigma-Aldrich)). Medium was changed every three days. Calcium imaging of DM-derived skeletal muscle activity in GCaMP reporter line (Gen1C)^37^ was performed using Nikon A1R MP (Multiphoton+N-STORM).

### *In vivo* xeno-transplantation of PSM derivatives

NOD/ShiJic-*scid*Jcl mice were purchased from CLEA Japan. Human sclerotome cells derived from healthy donor (WT) or homozygous/heterozygous luciferase reporter lines (625-A4 and 625-D4) were dissociated using Accutase (Life Tech) and resuspended in 100 µl of CDMi before being mixed with the same volume of Matrigel as previously described^4^. Numbers of transplanted cells ranged from ∼ 5.0 × 10^5^ – 1.2 × 10^6^ cells/injection. Cells were injected into mice subcutaneously with 26 G needle and 1 ml syringe (Terumo). Forming cartilage and bone tissues were taken out at 2 months post injection. Bioluminescence images were taken with IVIS Spectrum (PerkinElmer). Whole mount photos were taken with LEICA M205FA (Leica). Animal experiments were approved by the institutional animal committee of Kyoto University and performed in strict accordance with the Regulation on Animal Experimentation at Kyoto University.

### Quantitative real-time PCR (qRT-PCR)

RNA was extracted with the RNeasy mini kit (Qiagen) following the manufacturer’s instructions. cDNA was synthesized using Superscript III Reverse Transcriptase (Invitrogen) from 1 µg total RNA. cDNA was diluted 1:10 in RNase-free water. qRT-PCR was performed using Thunderbird SYBR qPCR Mix (Toyobo) and QuantStudio™ 12K Flex Real-Time PCR System (Thermo Fisher). The expression values of target genes were normalized by b-actin expression from the same cDNA templates. Details of utilized qRT-PCR primers are listed in Extended Data Table 5.

### Immunocytochemistry

Cells were fixed with 2% paraformaldehyde (PFA) for 30 minutes and washed twice with PBS. Samples were permeabilized with 0.2% Triton^®^ X-100 (Sigma-Aldrich) in PBS for 10 minutes at room temperature and then washed with PBST (1% TWEEN^®^ 20 (Sigma-Aldrich) in PBS). Subsequently, samples were blocked in 5% skim milk for 1 hour at room temperature and then stained with primary antibodies for overnight at 4°C. Samples were then washed with PBST three times and stained with secondary antibodies for 1 hour at room temperature. Antibodies were diluted in 10% blocking solution (5% skim milk) in PBST, washed with PBST twice and stained with DAPI for nuclear counterstaining for 5 minutes at room temperature. All images were taken using Nikon A1R MP (Multiphoton+N-STORM). For details of used primary and secondary antibodies see Extended Data Table 6.

### Histological analysis

Tissues were fixed with 4% PFA overnight at 4°C. Fixed samples were washed with PBS twice and embedded in paraffin. Sections were sliced at 3 µm for immunostaining and 5 µm for other stainings. Sections were stained with Hematoxylin-Eosin (HE), Safranin O, von Kossa, Pentachrome, anti-type I Collagen (COL1) antibody, anti-type II Collagen (COL2) antibody, and anti-Human Nuclear Antigen (HNA) antibody. Sections stained with antibodies were incubated for overnight at 4°C. Secondary antibodies were applied with N-Histofine^®^ Simple Stain™ MAX PO (Nichirei Bioscience Inc.) for 30 minutes at room temperature. Signals were detected by N-Histofine^®^ DAB-3S kit (Nichirei Bioscience Inc.). Details of used antibodies are listed in Extended Data Table 6.

### Flow cytometric analysis

Cells were washed with PBS and dissociated using Accutase (Life Technologies) and centrifuged. Cells were resuspended (1.0 × 10^7^ cells/ml) in FACS buffer (0.1% BSA in PBS) and stained with allophycocyanin (APC)-conjugated DLL1 antibody for 30 minutes at 4°C. Then, cells were stained with DAPI to eliminate dead cells after washing with FACS buffer once and finally strained through a filter mesh. As for the co-staining of intracellular molecules TBX6 and BRACHYURY with DLL1, cells were fixed with 4% paraformaldehyde (PFA) for 20 minutes at 4°C after initial staining with DLL1 antibody and washed twice with staining medium, which contained PBS with 2% fetal bovine serum (FBS). Samples were permeabilized with BD Perm/Wash buffer (BD Biosciences) for 15 minutes at room temperature and stained with TBX6 primary antibody or phycoerythrin (PE)-conjugated BRACHYURY antibody for 60 minutes at room temperature and washed with BD Perm/Wash buffer twice. The cells stained with TBX6 antibody were stained with Alexa Fluor^®^ 488-conjugated secondary antibody for 60 minutes at room temperature. The samples were washed with BD Perm/Wash buffer twice and suspended into staining medium. Flow cytometric analysis was performed using LSR or BD FACSAria II cell sorter (BD Biosciences). FACS data was analyzed and graphs were generated using FlowJo software (FlowJo LLC). For details of used antibodies see Extended Data Table 6.

### Reporter constructs

For the HES7-reporter, human *HES7* promoter (5937 bp) and 3’UTR were fused to Luciferase2-NLS-d1PEST^38^. For the dual reporter assay, the *HES7* promoter and 3’UTR were fused to NanoLuc-NLS-d1PEST, while human *DKK1* promoter (2218 bp) and 3’UTR were fused to Luciferase2-NLS-d1PEST. These reporters were integrated into the genome using *piggyBac* transposition. See Fig. 1g and Fig. 2d for schematic overviews of used reporter constructs.

### 2D-oscillation assay

Luminescence was measured in the presence of D-luciferin (200 µM) with Kronos Dio Luminometer (Atto). For the dual reporter assay, HES7- and DKK1-reporter constructs were simultaneously introduced into the cells, and each luminescence was filtered and measured in the presence of Furimazine (400 nM) and D-luciferin (1 mM). HES7-reporter cells were seeded on a 35 mm dish coated with iMatrix-511 at 3000 cells/dish. After 4 days culture, medium was changed into CDMi containing SB431542 (10 µM), DMH1 (2 µM), CHIR99021 (10 µM) and bFGF (20 ng/ml). After additional 3 days culture, the medium was changed into CDMi without inhibitors for measurement with Kronos Dio Luminometer (Atto). This (modified) one-step protocol^39^ was used for Fig. 1g and 2d. All other oscillation measurements were performed using our standard step-wise PSM induction protocol.

### 3D-synchronization assay

To make 3D cell spheres, HES7-reporter iPSCs were seeded into non-adhesive round bottom 96 well plates at 1000-3000 cells/well and cultured in CDMi containing BMP4 (50 ng/ml), CHIR99021 (10 µM) and Y27632 (10 µM). After one day of culture, Y27632 was removed. After 18 hours culture the medium was changed into CDMi containing DMH1 (2 µM) and CHIR99021 (10 µM). After 6 hours culture, the cell sphere was transferred to a fibronectin-coated glass bottom dish with CDMi containing DMH1 (2 µM) and D-luciferin (1 mM), and luminescence was imaged with a customized incubator microscope LCV110 (Olympus). Obtained Image was analyzed with Metamorph (Molecular Devices) and Excel (Microsoft), and kymograph was made by averaging signals over 10 pixels with Metamorph.

### Sampling for RNA-seq analysis of oscillating genes

Our standard step-wise PSM induction protocol was used with the following modifications. HES7-reporter cells were seeded on a 35 mm dish coated with Matrigel. At 12 hours during the 2nd-step (PSM induction), the cells were split into multiple 35 mm dishes at 4.0 × 10^5^ cells/dish and cultured in CDMi containing SB431542 (10 µM), LDN193189 (250 nM) and CHIR99021 (3 µM). After 12 hours culture the medium was changed into CDMi containing SB431542 (10 µM), LDN193189 (250 nM), CHIR99021 (3 µM) and bFGF (20 ng/ml). The luminescence was continuously monitored with Kronos Dio Luminometer with one sample, and the other samples were frozen at each time point.

### Library preparation for RNA-sequencing

Total RNA was extracted using RNeasy mini kit (Qiagen) following the manufacturer’s instructions. RNA-seq libraries were generated from 200-300 ng total RNA with the TruSeq Stranded mRNA LT Sample Prep Kit (Illumina, San Diego, CA, USA) according to the manufacturer’s protocol, with the exception of the libraries used for RNAseq analysis of oscillating genes, which were generated from 120 ng total RNA using NeoPrep system (Illumina) following the manufacturer’s instructions. The obtained RNA-seq libraries were sequenced on NextSeq 500 (75 bp – 86 bp single-end reads, Illumina).

### RNA-sequencing data analyses

The sequenced reads were mapped to the hg38 human reference genome plus the luciferase reporter sequence using HISAT2 (version 2.1.0)^40^ with the GENCODE v25 annotation gtf file after trimming adaptor sequences and low-quality bases by cutadapt-1.14^41^. The mapped reads with high mapping quality (MAPQ >= 20) were used for further analyses. For identification of the differentiation stage-related genes, the differentially expressed genes (>= 5-folds changes and q-values =< 0.05 between any pair of samples) were extracted using Cuffdiff^42^ within Cufflinks version 2.2.1 package and GENCODE v25 annotation file, and the extracted genes were grouped into six stages based on the maximum expression levels (FPKM values determined by Cuffdiff) among the differentiation stages. The low expressed genes (=< 10 FPKM) across all stages were filtered out before grouping. For identification of the oscillation genes, the uniquely mapped reads were counted and normalized to calculate the gene expression levels using HTSeq (version 0.6.1)^43^ with GENCODE v25 annotation gtf file (protein-coding genes) and edgeR (version 3.18.1)^44^ after filtering low expressed genes (cpm =< 1) across all conditions in each experiment, while rhythmic genes were identified by ARSER (version 2.0)^45^ with FDR_BH =< 0.03 in both two independent experiments. The filtering genes for noise judged by ARSER in both experiments were excluded from oscillation genes. For pathway and Gene Ontology analyses, DAVID web tools^46^ and IPA (Qiagen Inc., https://www.qiagenbioinformatics.com/products/ingenuitypathway-analysis) were used. For identification of the patient (STD-A/F) related genes, fold changes with q-values were calculated with HTSeq (version 0.6.1)^43^ with GENCODE v25 annotation gtf file and edgeR (version 3.18.1)^44^. The genes whose expression values were upregulated or downregulated (>= 3 folds changes, q<0.05, STD lines vs. WT), and increases or decreases were inhibited (>=50%, q<0.05) in the rescued lines, were defined as STD-related upregulated or downregulated genes, respectively. The genes whose expression levels were low (average cpm =< 5) both in wild type and STD-lines were filtered out. For comparisons of expression profiles between knock-out (KO) cell lines and their parental cell lines (wild type), FPKM values, fold changes and q-values were calculated using Cuffdiff^42^ within Cufflinks version 2.2.1 package and GENCODE v25 annotation file (protein-coding genes). The principal component analysis (PCA) and representation of heatmaps and scatter plots were performed using R software.

### CRISPR/Cas9 gene knock-out

Gene knock-out was performed using transient transfection of pSpCas9(BB)-2A-Puro (PX459) V2.0 (a gift from Feng Zhang, Addgene plasmid #62988). Oligonucleotides encoding sgRNA protospacer sequences (Extended Data Table 5) were annealed and cloned as described previously^47^. sgRNAs were verified by sequencing. Plasmid DNA (1 ug) was transfected into iPS cells by electroporation followed by selection with 0.5 µg/mL puromycin for 48 hours. Surviving cells were allowed to recover and then replated at low density before picking isolated colonies. For overview of knock-out reporter line establishment and details of used sgRNAs see Extended Data Fig. 4 and Extended Data Table 5.4.

### Whole exome sequencing and variant calling

Whole exome sequencing was performed as previously described^48,49^. Briefly, DNA (3 μg) was sheared with S2 Focused-ultrasonicator (Covaris, Wobum, MA, USA) and processed by SureSelectXT Human All Exon V5 (Agilent Technologies, Santa Clara, CA, USA). Captured DNA was sequenced using HiSeq 2000 (Illumina, San Diego, CA, USA) with 101 bp pair-end reads with seven indices. Image analysis and base calling were performed using HCS, RTA and CASAVA software (Illumina). Reads were mapped to the reference human genome (hg19) by Novoalign-3.02.04. Aligned reads were processed by Picard to remove PCR duplicates. Variants were called by GATK v2.7-4 following GATK Best Practice Workflow v3^50^ and annotated by ANNOVAR^51^. All the variants of the candidate genes, which have been reported to cause SCD or CS, were evaluated using five databases: gnomAD, Human Gene Mutation Database (HGMD), SIFT, PolyPhen-2, MutationTaster.

### Quality control of established iPSCs

Morphological images of iPSC colonies were captured using Olympus CKX41 microscope with a PlanApo 10×/0.75 objective lens (Olympus) and Nikon digital camera DS-Fil. Chromosomal G-banding analyses were performed by Chromocenter Inc, Tottori, Japan. Genomic DNA and total RNA were extracted with AllPrep DNA/RNA mini kit (Qiagen) following the manufacturer’s instructions. Genomic DNA was diluted into 25 ng/ml in distilled water. cDNA was synthesized using PrimeScript™ RT Master Mix (Takara) from 500 ng total RNA and diluted 1:10 in RNase-free water for *OCT3/4* and *NANOG* mRNA expression analysis, and 1 μg total RNA for TaqMan™ hPSC Scorecard™ analysis. *OCT3/4* and *NANOG* mRNA expression were confirmed by quantitative real-time PCR (qRT-PCR) with TaqMan™ assay using StepOnePlus™ Real-Time PCR Systems (Thermo Fisher). Primers and probe sequences are provided in Extended Data Table 5.2. The expression values of target genes were normalized by *GAPDH* expression from the same cDNA templates and calculated relative to 201B7 iPS cell line. Residual plasmids used for iPSC establishment were analyzed by TaqMan™ quantitative PCR using StepOnePlus™ Real-Time PCR Systems (Thermo Fisher). Primer and probe sequences (cmCAG: common-CAG) are designed on CAG-promoter region included in all of the episomal vectors for iPSC generation and listed in Extended Data Table 5.2. The residual plasmid numbers were determined by a standard curve method with pCE-OCT3/4 episomal plasmid of known quantity using 50 ng genomic DNA of STD-iPSCs at passage 6.

### Initial validation of established iPSCs

Established (patient) iPSCs together with control human PSCs were differentiated into ectoderm, mesoderm and endoderm lineages using STEMdiff™ Trilineage Differentiation Kit (STEMCELL Technologies). hPSCs reaching 70–80% confluency were harvested with TrypLE™ Select Enzyme (1X) (Thermo Fisher) and plated as a single cell suspension in mTeSR1 medium (STEMCELL Technologies) containing 10 μM Y27632 (Wako) on 6-well plates coated with Matrigel (BD Biosciences). The cells were plated at 4.0 × 10^5^ cells, 2.0 × 10^5^ cells and 4.0 × 10^5^ cells per well for ectoderm, mesoderm and endoderm differentiation culture respectively and differentiated following the manufacturer’s instructions. For FACS-based evaluation of undifferentiated PSCs and each of the three germ layers (1.0 × 10^6^ cells each) were fixed with 4% paraformaldehyde phosphate buffer solution (4% PFA/PBS) for 20 minutes at 4°C and washed twice with staining medium, which contained PBS with 2% fetal bovine serum (FBS). Samples were permeabilized with BD Perm/Wash buffer (BD Biosciences) for 15 minutes at room temperature and stained with fluorescence-conjugated primary antibodies listed in Extended Data Table 6.3. The samples were washed with BD Perm/Wash buffer twice and suspended into staining medium. Flow cytometric analysis was performed using LSR (BD Biosciences). FACS data was analyzed and graphs were generated using FlowJo software (FlowJo LLC). For transcript level assessment of differentiation capacity, qPCR was performed with 384-well TaqMan™ hPSC Scorecard™ panel (Thermo Fisher) by QuantStudio™ 12K Flex Real-Time PCR System (Thermo Fisher) using undifferentiated PSC and each of the three germ layers cDNA samples. Pluripotency and differentiation property into ectoderm, mesoderm and endoderm lineages were scored by hPSC Scorecard Analysis software, which is available on the Thermo Fisher website (https://www.thermofisher.com/jp/en/home/life-science/stem-cell-research/taqman-hpsc-scorecard-panel.html).

### Gene correction of patient iPSCs

Correction of mutations in patient iPSCs was performed using the MhAX method as previously described^28^. Briefly, donor plasmids for correction of each mutant allele were created by PCR amplification of the right arm from the cloned genomic DNA of STD patient cells corresponding to the matching mutant allele, and the left arm from normal 201B7 iPSC using the common primers listed in Extended Data Table 5. InFusion cloning (Clontech) was used to assemble the arms with a restriction-digested CAG::mCherry-IRES-puro selection cassette (Addgene plasmid #113876) and CAG::GFP plasmid backbone (Addgene plasmid #107281). PCR-amplified regions and InFusion junctions were verified by sequencing. Oligonucleotides encoding sgRNA protospacer sequences (Extended Data Table 5) were annealed and cloned into pX330-U6-Chimeric_BB-CBh-hSpCas9 (a gift from Feng Zhang, Addgene plasmid # 42230) as described previously^47^. sgRNAs were verified by sequencing. For gene targeting, allele-matched donor plasmids (3 µg) and Cas9 / sgRNA expression plasmids (1 µg) were co-transfected by electroporation into 1.0 × 10^6^ STD-A/F patient iPSCs, which were then divided and plated under feeder-free conditions for 48 hours in AK02N medium (Ajinomoto) containing 10 µM ROCK inhibitor Y27632 (Wako) before initiating antibiotic selection (0.5 µg/mL puromycin, Sigma-Aldrich). Nine days after plating, puromycin-resistant cells were pooled and passaged. GFP^neg^ / mCh^pos^ colonies were isolated, cultured, stored, and processed for genomic DNA isolation under feeder-free conditions in 96-well format. iPSC clones positive for PCR genotyping and sequencing were defrosted and expanded for genomic DNA extraction and Southern blot verification. For cassette excision, 3 µg of the pX-EGFP-g1 expression plasmid (Addgene plasmid #107273)^28^ was transfected into 1.0 × 10^6^ gene-targeted patient iPSCs, which were then divided and plated under feeder-free conditions for 48 hours in AK02N medium containing 10 μM Y27632, followed by growth without selection for a total of 6 days. mCh^neg^ cells were isolated by FACS on a BD FACSAria II cell sorter, and plated at low density for clonal isolation after 8 days. Isolated clones were cultured, stored in 96-well format, then genotyped for cassette excision by PCR and sequencing before final verification by Southern blot.

### Genomic DNA extraction

Genomic DNA for PCR amplification and sequencing was isolated from 0.5 – 1.0 × 10^6^ iPS cells using a DNeasy Blood and Tissue Kit (Qiagen). Genomic DNA for Southern blotting was isolated from a single confluent well of a 6-well dish using lysis buffer (100 mM Tris-HCl, pH 8.5, 5 mM EDTA, 0.2% SDS, 200 mM NaCl, and 1 mg/mL Proteinase K) followed by phenol-chloroform extraction and ethanol precipitation from the aqueous phase. Genomic DNA was eluted from columns or resuspended from precipitate in TE pH 8.0.

### iPSC genotyping

PCR primers flanking annotated coding exons of *DLL3* (Accession NG_008256.1), *HES7* (Accession NG_015816.1), and *LFNG* (Accession NG_008109.2), *MESP2* (Accession NG_008608.1) were designed using NCBI Primer-BLAST with optional settings filtering human repeats and SNPs, with primer pair specificity checking to H. sapiens (taxid:9606). PCR primers for genotyping gene-edited cell lines were designed using similar principles. All genotyping primers are listed in Extended Data Table 5. Genomic PCR was carried out using KAPA HiFi HotStart (KAPA Biosystems) on a Veriti 96-well Thermal Cycler (Applied Biosystems) according to the manufacturer’s instructions. Specific PCR conditions are available upon request. PCR products were treated with ExoSAP-IT Express (Affymetrix) and sequenced with the primers indicated in Extended Data Table 5 using BigDye Terminator v3.1 Cycle Sequencing Kit (Applied Biosystems) on a 3130xl Genetic Analyzer (Applied Biosystems). Sequence analysis was performed using variant calling in Sequencher (Genecodes) or alignment in Snapgene (GSL Biotech LLC.). For *MESP2* gene correction, patient and rescued iPS cells were analyzed by Southern blotting as described previously^28^. Probe regions were PCR amplified with Ex Taq (Takara) directly from genomic DNA or cloned plasmid templates to incorporate DIG-labeled dUTP (Roche) using the primers described in Extended Data Table 5. Genomic DNA (5-10 μg) was digested with *Eco*RI. Genomic DNA from patient iPSCs and iPSC clones rescued by gene editing were genotyped using an Infinium OmniExpress-24 v1.2 (Illumina) SNP array according to the manufacturer’s recommendations. Data collection was performed on an iScan Bead Array Scanner (Illumina). Data was compared to the reference human genome (hg19) using a combination of PennCNV, cnvPartition, GWAS tools, and MAD. Karyograms were prepared in R (version 3.2.5) using GWASTools (version 1.16.1)^52^.

### Data accessibility

All RNA sequencing data utilized for this study have been deposited in Gene Expression Omnibus (https://www.ncbi.nlm.nih.gov/geo/) under the accession number GSE116935.

## EXTENDED DATA FIGURES

**Extended Data Fig. 1.**
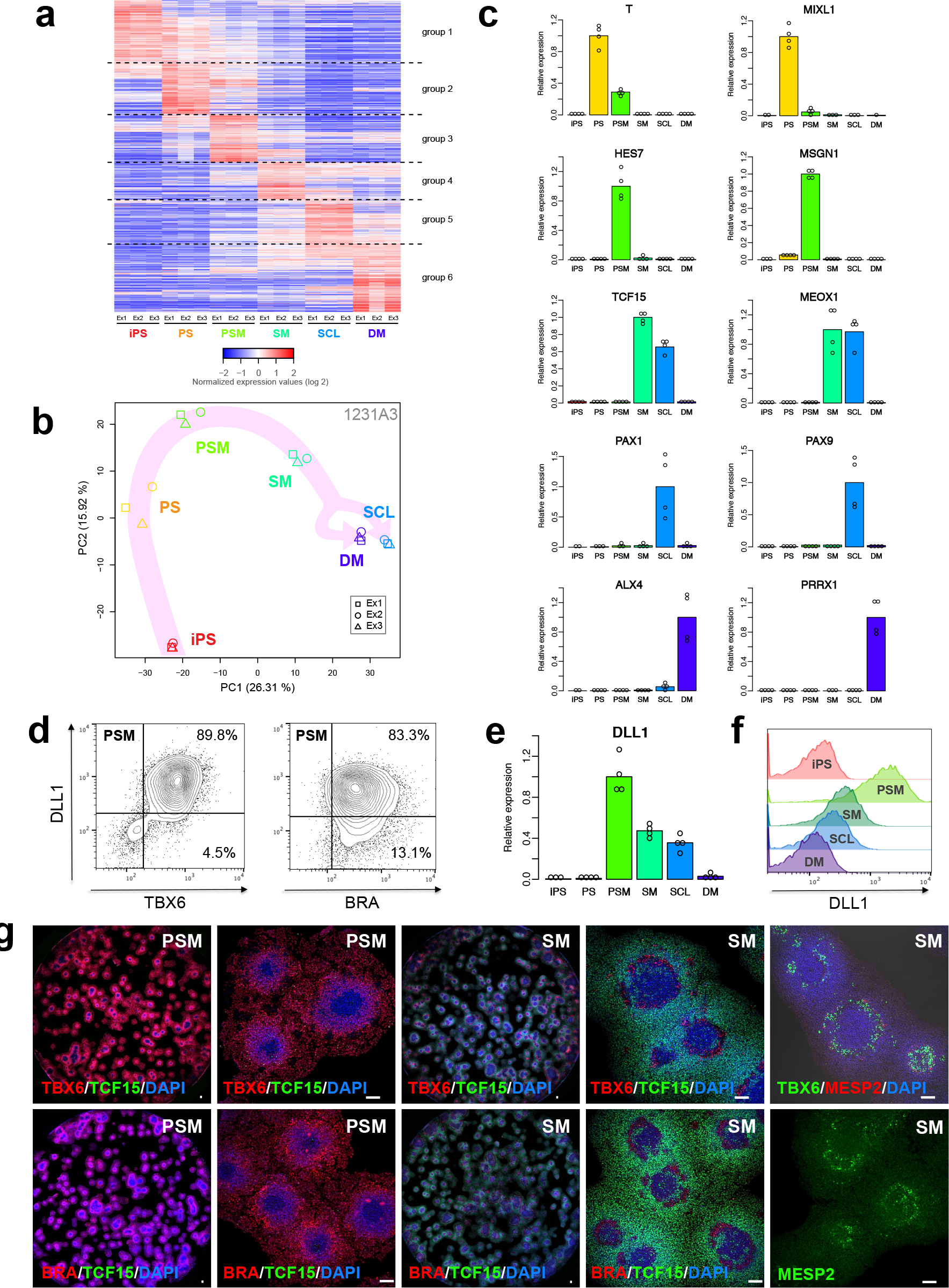
Characterization of step-wise induced human presomitic (PSM) and somitic mesoderm (SM). **a**, Heatmap of gene expression levels in step-wise induced human PSM and its derivatives (using iPSC-line 1231A3). FPKM values of each gene were normalized to mean of all samples. The gene order is the same as in Fig. 1b. **b**, PCA analysis of transcript expression levels in human PSM and derivatives (1231A3). **c**, qRT-PCR based validation of RNA-seq results; summary of four independent experiments with three technical replicas each using 201B7. Similar results were obtained for 1231A3 (data not shown). It should be noted that open circles in some conditions are less than four because no Ct values in the samples were obtained after 45 cycles of PCR to calculate expression values. **d**, Flow cytometry-based evaluation of DLL1 and TBX6 (left) as well as DLL1 and BRACHYURY (BRA) (right) expression at PSM stage (1231A3). **e**, Expression of *DLL1* on transcript level throughout the different stages of induction (201B7). **f**, Expression of DLL1 on protein level. Correlation of FACS data with qRT-PCR results (201B7) shown in (e). **g**, Immunofluorescence staining of PSM markers TBX6 and BRA as well as somitic mesoderm marker TCF15 at PSM (left half panels) and SM (right half panels) stages; entire wells (left) and magnified views of selected areas (right). Staining of segmentation marker MESP2 (alone or co-staining with TBX6) shown in far right side of panel. Scale bar: 100 μm.

**Extended Data Fig. 2.**
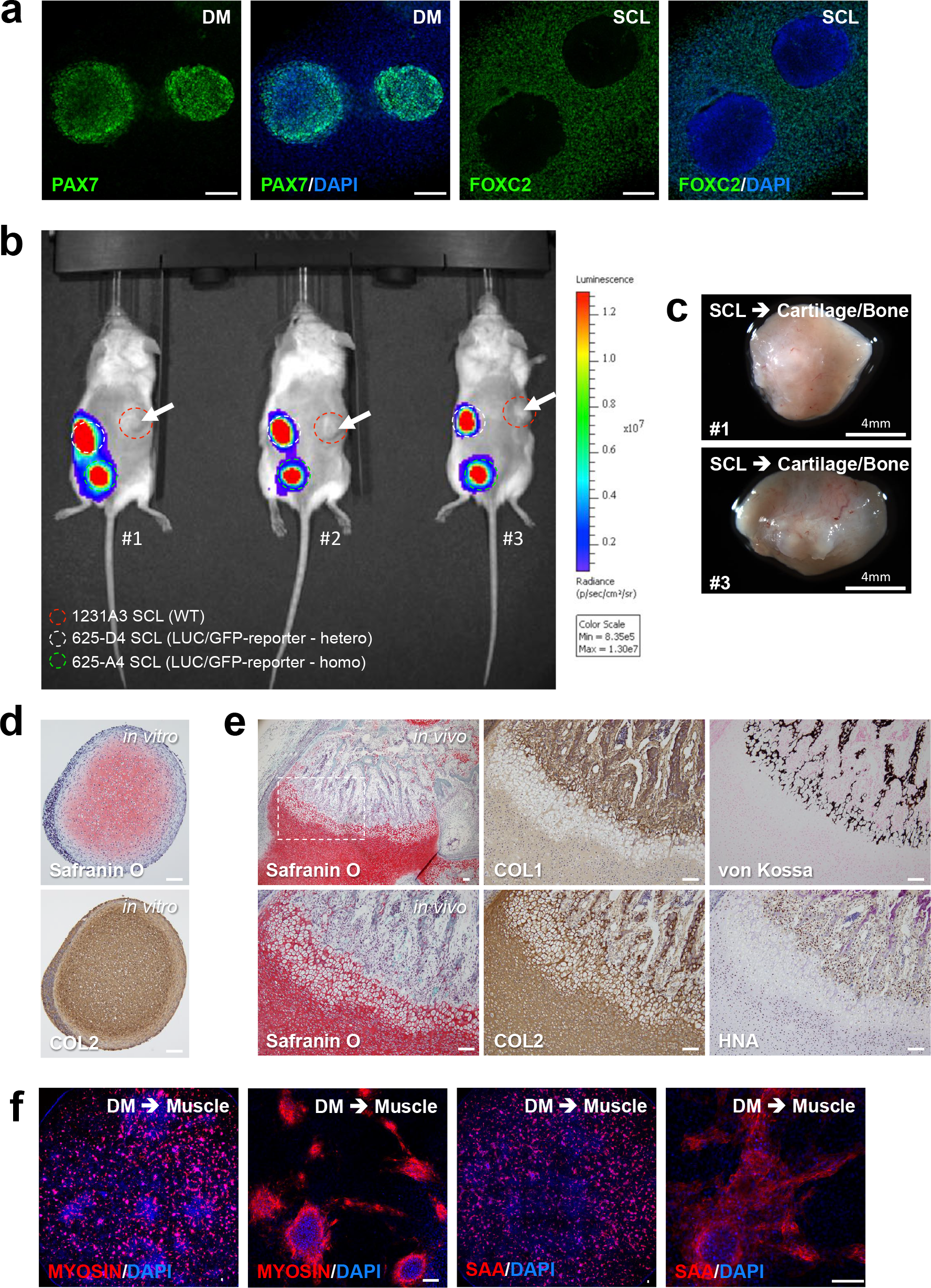
Molecular and functional characterization of step-wise induced human PSM-derivatives, sclerotome (SCL) and dermomyotome (DM). **a**, Developmental-stage specific expression of DM (PAX7) and SCL (FOXC2) markers on protein level (201B7). Scale bar: 100 μm. **b**, Assessment of *in vivo* bone and cartilage forming ability of human sclerotome (SCL). Subcutaneous transplantation of PSC-derived SCL step-wise induced from WT (1231A3) and luciferase reporter lines (625-D4 and 625-A4). Evaluation of transplanted cells via IVIS at two months post-transplantation. Injection sides are marked by dashed/colored circles. Cartilage and bone forming areas of WT iPSC line (1231A3) marked by white arrows. **c**, Whole-mount images of WT SCL-derived *in vivo* cartilage/bone tissues isolated from transplanted mice #1 and #3. Explant isolated from mice #2 is shown in Fig. 1i. **d**, Staining of *in vitro* human SCL derived cartilage (3D-CI) sections. Observed Safranin O and type II collagen (COL2) signals are indicative of *in vitro* cartilage formation. **e**, Sections and staining of area shown in Fig. 1i; Safranin O and COL2 staining in human *in vivo* SCL-derived cartilage areas; von Kossa and COL1 staining in ossifying cartilage and forming bone areas. Majority of cells contributing to cartilage/bone formation are HNA positive and of human origin (right lower panel). Scale bar: 100 μm. **f**, Evaluation of *in vitro* muscle induction from human DM. Myosin and sarcomeric alpha-actinin (SAA) staining of *in vitro* DM-derived skeletal muscle; representative images of entire well (left) and magnified areas (right). Scale bar: 100 μm.

**Extended Data Fig. 3.**
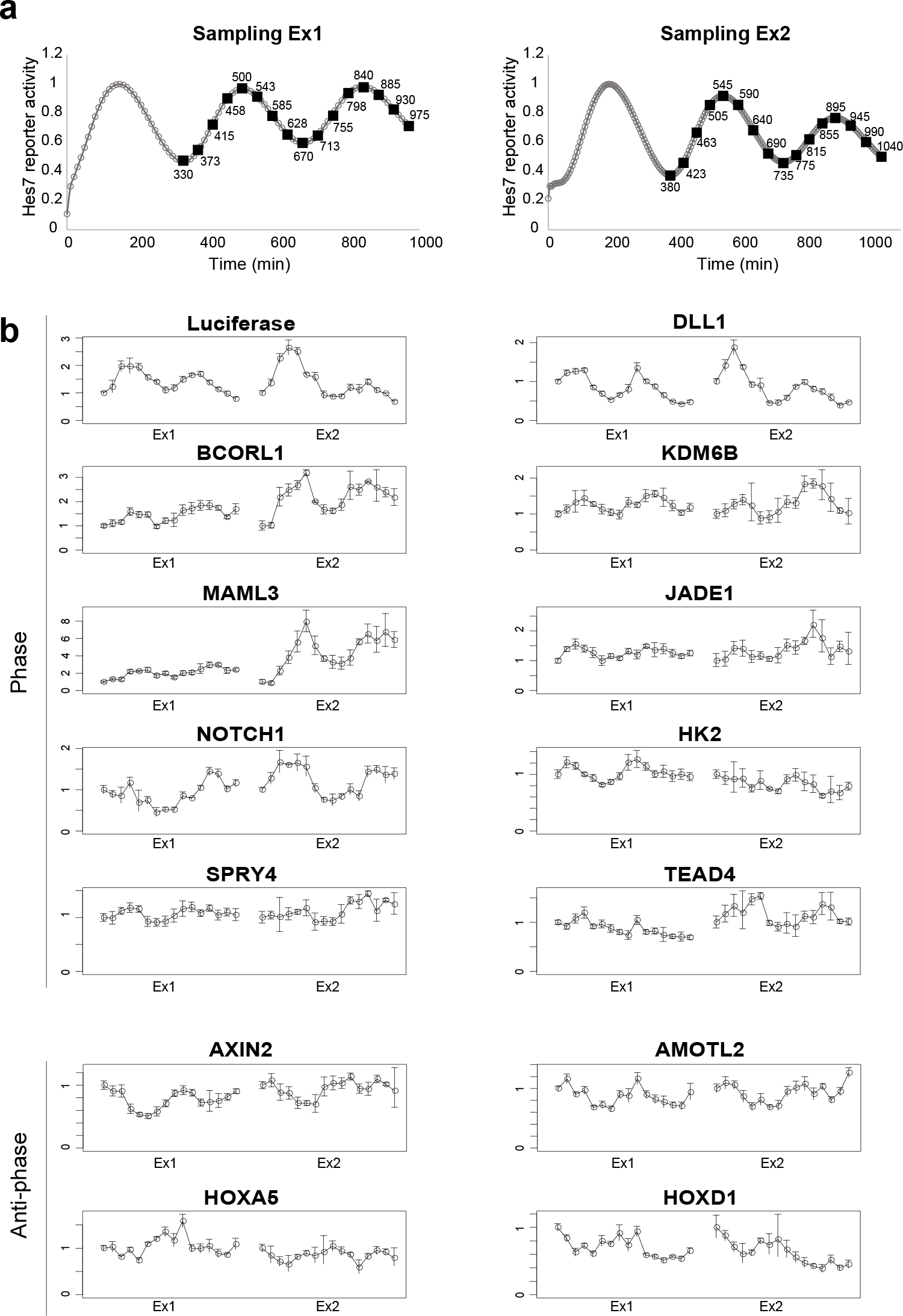
Isolation and transcriptome analysis of timely coordinated human PSM samples. **a**, Sampling for RNA-seq. HES7-reporter activity was continuously monitored with one sample, and the other samples were frozen at each time point indicated in the graph. **b**, qRT-PCR validation of identified novel phase and anti-phase oscillating genes. *DLL1* and *AXIN2* show clear phase or anti-phase oscillation despite not being included into high stringency cut-off RNA-seq candidate list. Error bars indicate S.D. of three technical replicas for each time point and sample set.

**Extended Data Fig. 4.**
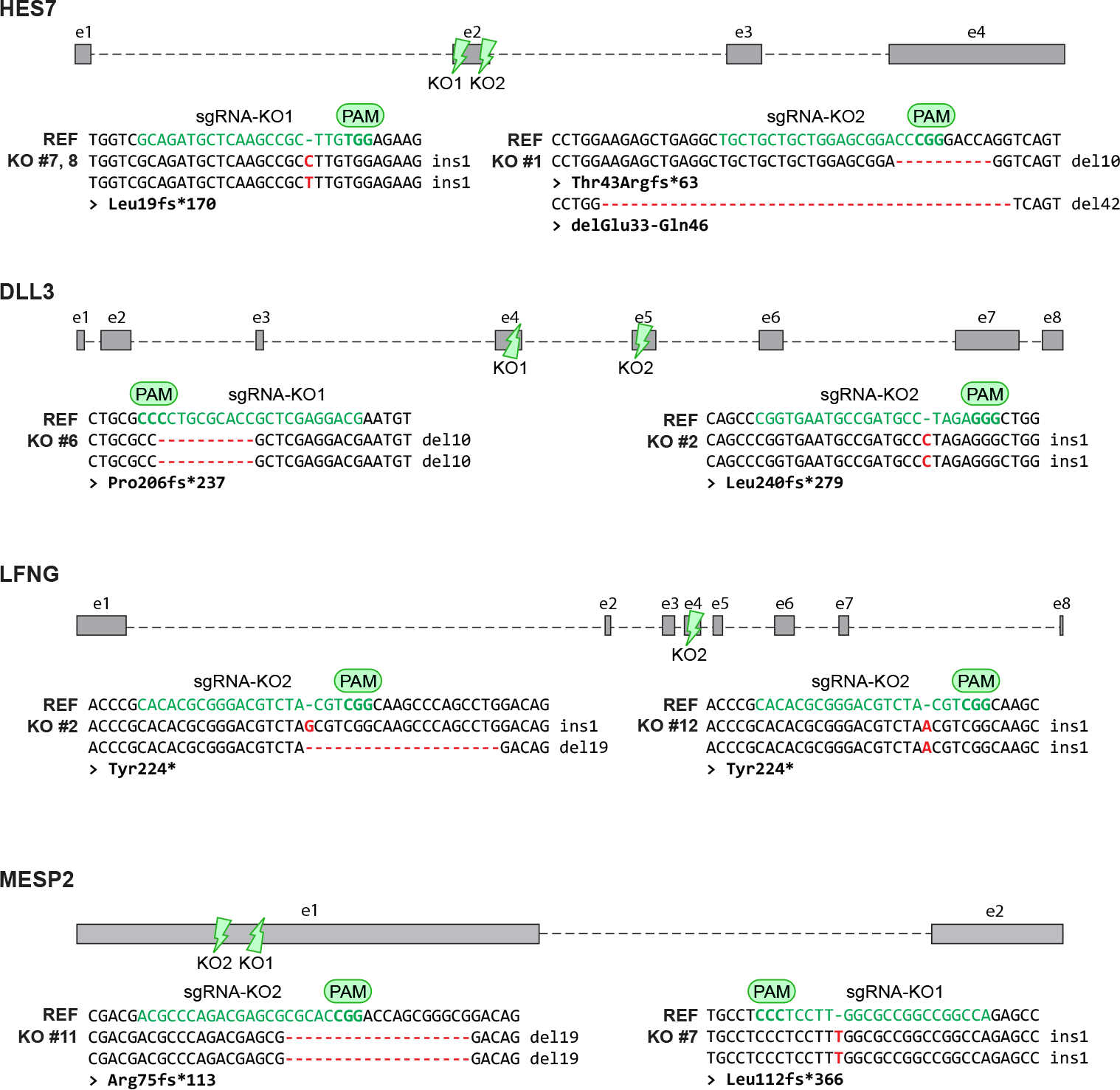
Overview of knock-out reporter line generation. Schematics of the *HES7*, *DLL3*, *LFNG*, and *MESP2* genes. Positions of the sgRNAs used in this study are shown. sgRNAs were designed to target at or near regions of known pathogenic mutations, particularly those resulting in frameshifts and premature termination. Sequence analysis of iPSC clones used in this study indicating indel mutations generated by Cas9. Predicted effects on the protein sequence are listed below the sequence alignments.

**Extended Data Fig. 5.**
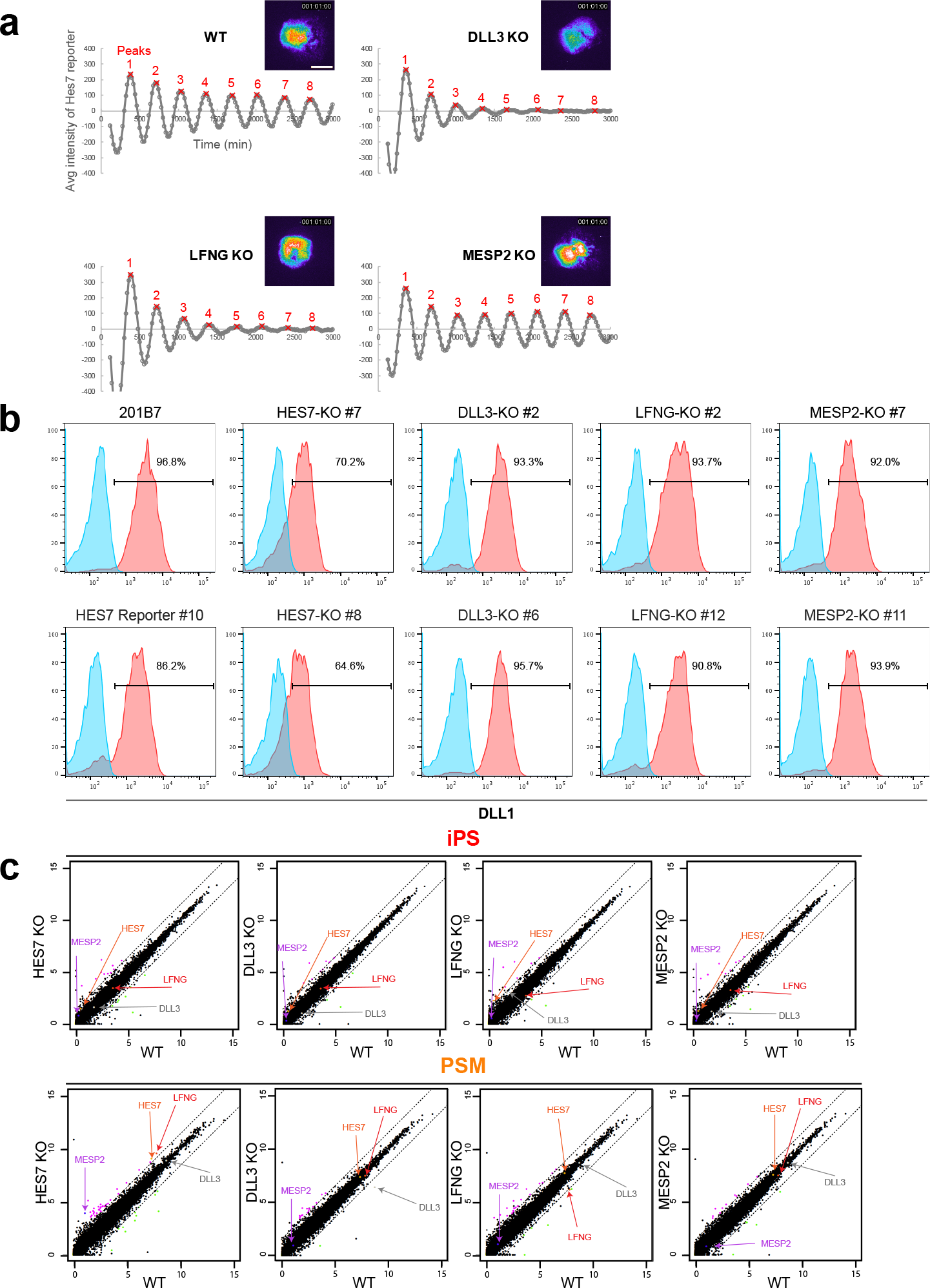
Characterization of knock-out reporter lines. **a**, Oscillation damping rate of knock-out PSMs in 3D-synchronization assay. Average signal of the entire image at each time point was measured using the sample shown in Extended Data Movie 3. The signal was detrended (± 2 hours), and each oscillation peak was detected to define the amplitude. Representative images and graphs of three independent experiments are shown. Scale bar: 500 μm. **b**, Evaluation of DLL1 expression via FACS analysis at iPSC and PSM stages of healthy control and knock-out iPSC lines. PSM induction efficiency is high in all analyzed samples; slight reduction of DLL1 induction efficiency in HES7-KO lines. Representative results of two different knock-out lines each are shown. **c**, Scatter plot of transcriptome analysis of wild type and KO lines at iPSC and PSM stages. Positions of expression values for *MESP2*, *DLL3*, *LFNG* and *HES7* are highlighted with colored arrows.

**Extended Data Fig. 6.**
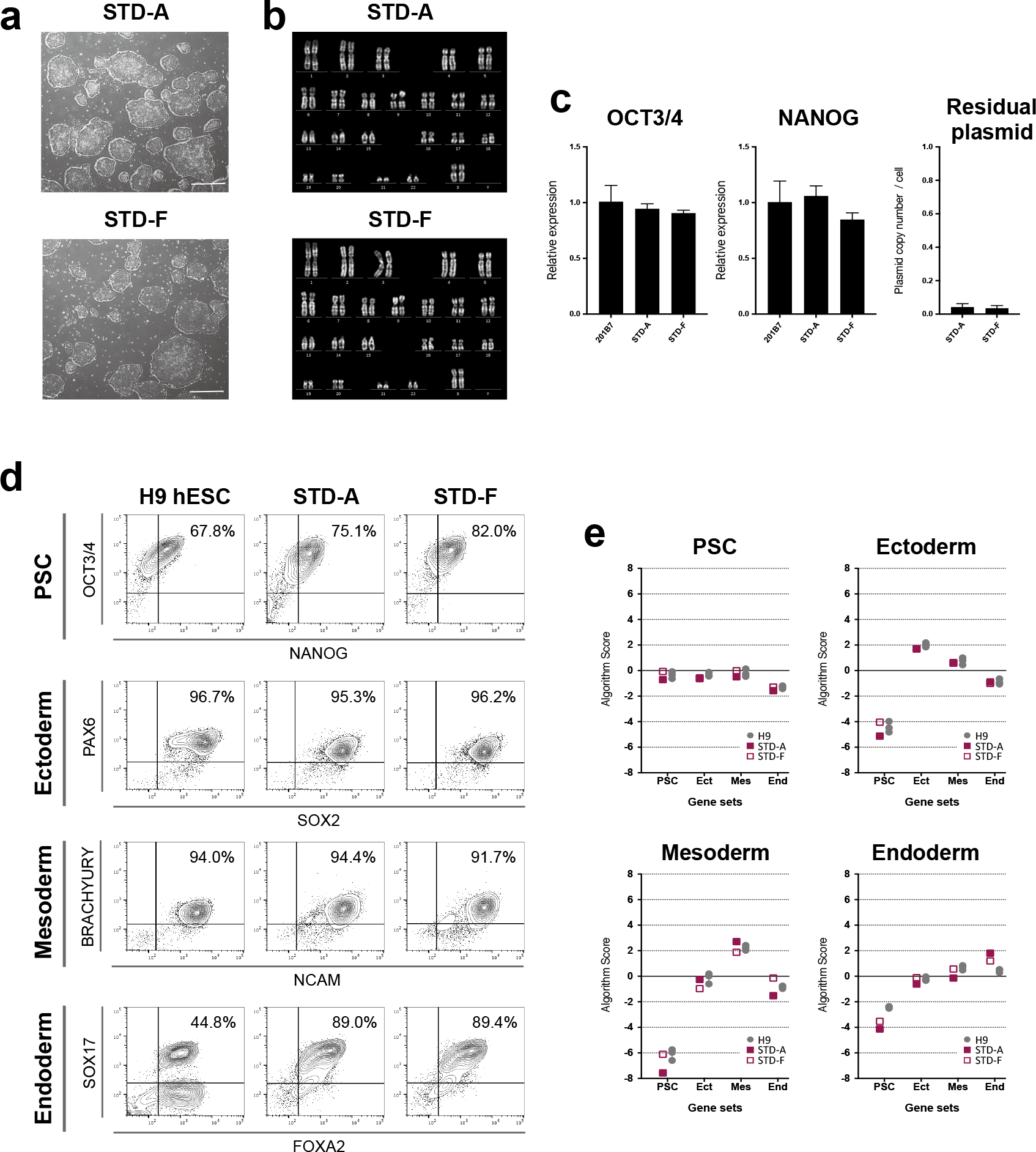
Initial characterization of STD patient iPSC line. **a**, Bright field view of STD iPSC clones STD-A and STD-F. Scale bar: 500 μm. **b**, Normal karyotype (46, XX) in both clones of STD patient iPSC line by chromosomal G-banding analysis. **c**, Expression of pluripotency markers *OCT3/4* and *NANOG* in STD-A and STD-F cells compared to WT iPSC line (201B7). Quantification of residual plasmid levels in STD clones (right side of panel). **d**, FACS-based evaluation of differentiation capacity into three germlayers of healthy control (H9 hESC) and patient lines (STD-A and STD-F). **e**, Quantification of differentiation capacity into ectoderm, mesoderm and endoderm at transcript level.

**Extended Data Fig. 7.**
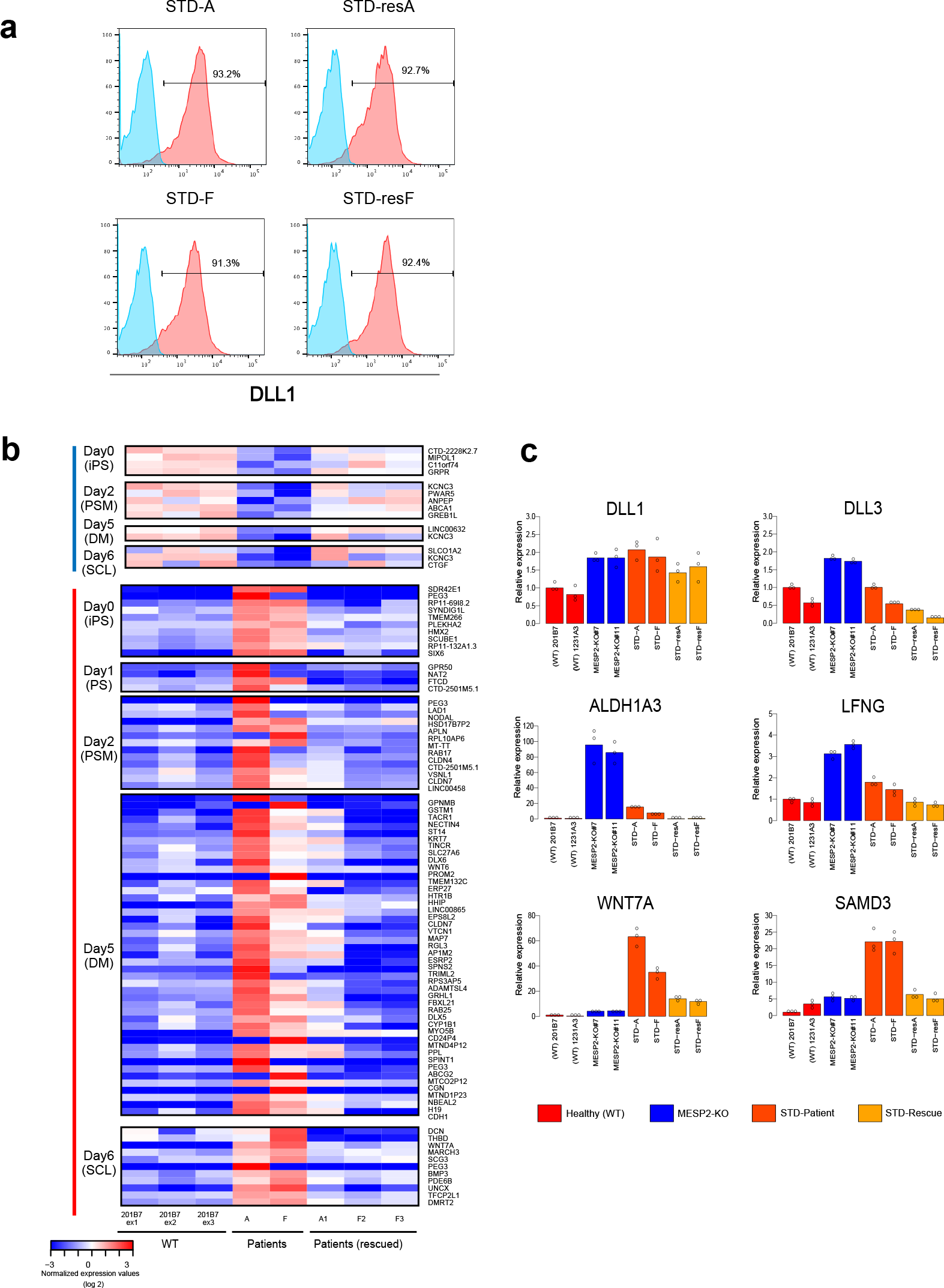
Comparison of patient vs. rescue STD lines. **a**, Representative DLL1 expression in PSM-derived from STD patient (STD-A and STD-F) and corresponding rescue lines (STD-resA and STD-resF); n=3. **b**, Heatmap of gene expression levels of transcripts differentially expressed in patient lines STD-A and STD-F, when compared to wild-type (201B7) and corrected rescue clones (STD-resA (A1) and STD-resF (F2 and F3)). Analysis covers all stages of step-wise PSM induction and differentiation. For SM stage data see Fig. 4e and Extended Fig. 7c. **c**, qRT-PCR-based validation of additional candidates found via RNA-seq to be up-regulated in STD patient lines at the SM stage. Data of three independent experiments are shown.

**Extended Data Fig. 8.**
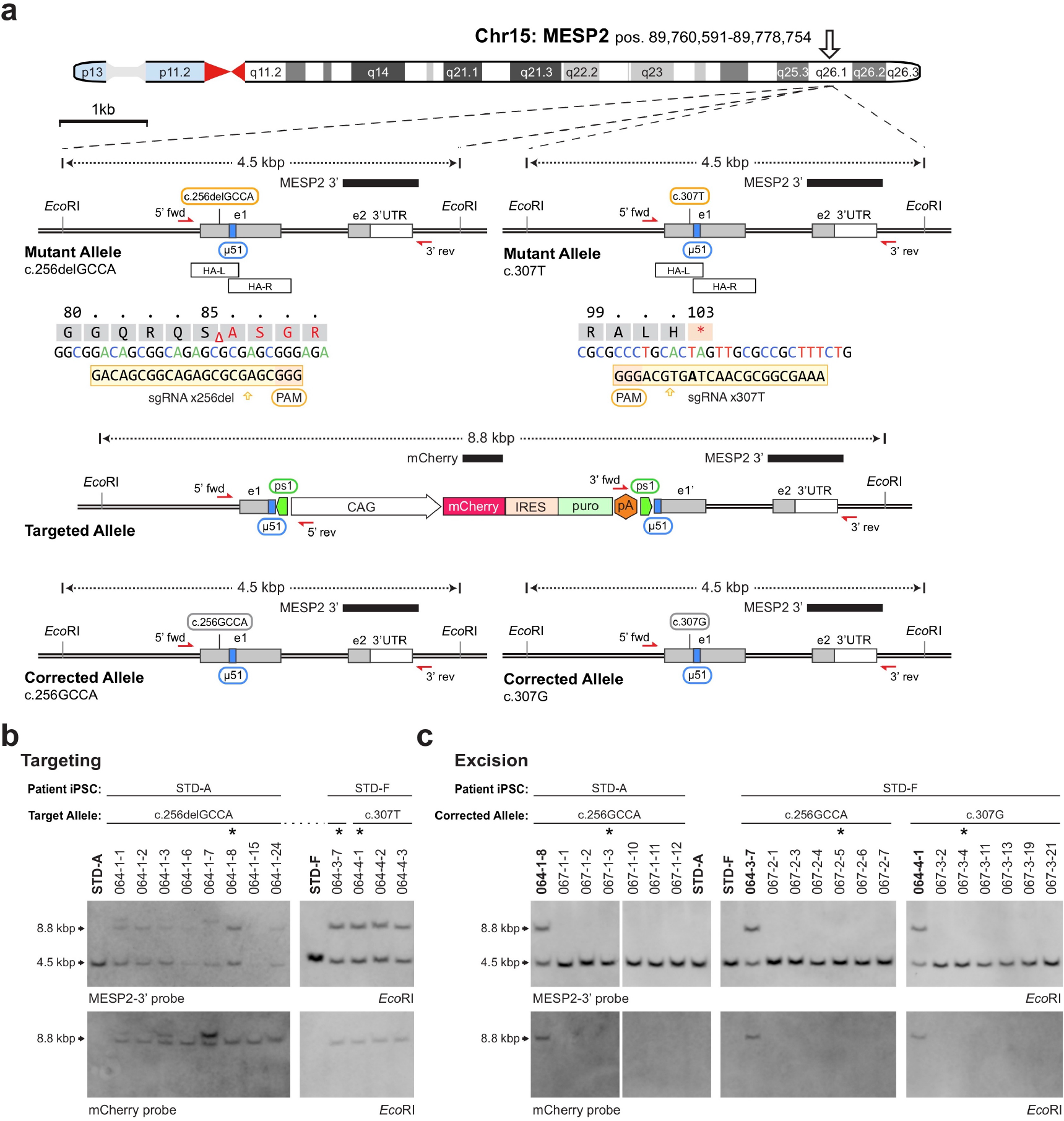
Allele-specific gene correction of *MESP2* in patient iPS cells. **a**, Detailed schematic of the gene correction strategy described in Fig. 4e, 4f. Depicted are the two mutant or corrected *MESP2* alleles with coding and non-coding exons (grey and white), overlapping donor vector homology arms (HA-L, HA-R), engineered 51 bp microhomology (μ51, blue), inverted protospacers for cassette excision (ps1, green), genotyping primers (red arrows), and Southern blotting probes (black bars). The sequences of mutation-specific sgRNAs are shown below each mutant allele. The gene-targeted intermediate shows details of the CAG::mCherry-IRES-puro cassette used for enrichment. **b**, Southern blot analysis of the targeted iPSC clones. Samples marked with an asterisk were selected for cassette excision. **c**, Southern blot analysis of gene corrected iPSC clones following selection marker removal. Samples marked with an asterisk were selected for phenotyping (067-1-3, STD-resA1; 067-2-5, STD-resF2; 067-3-4, STD-resF3).

**Extended Data Fig. 9.**
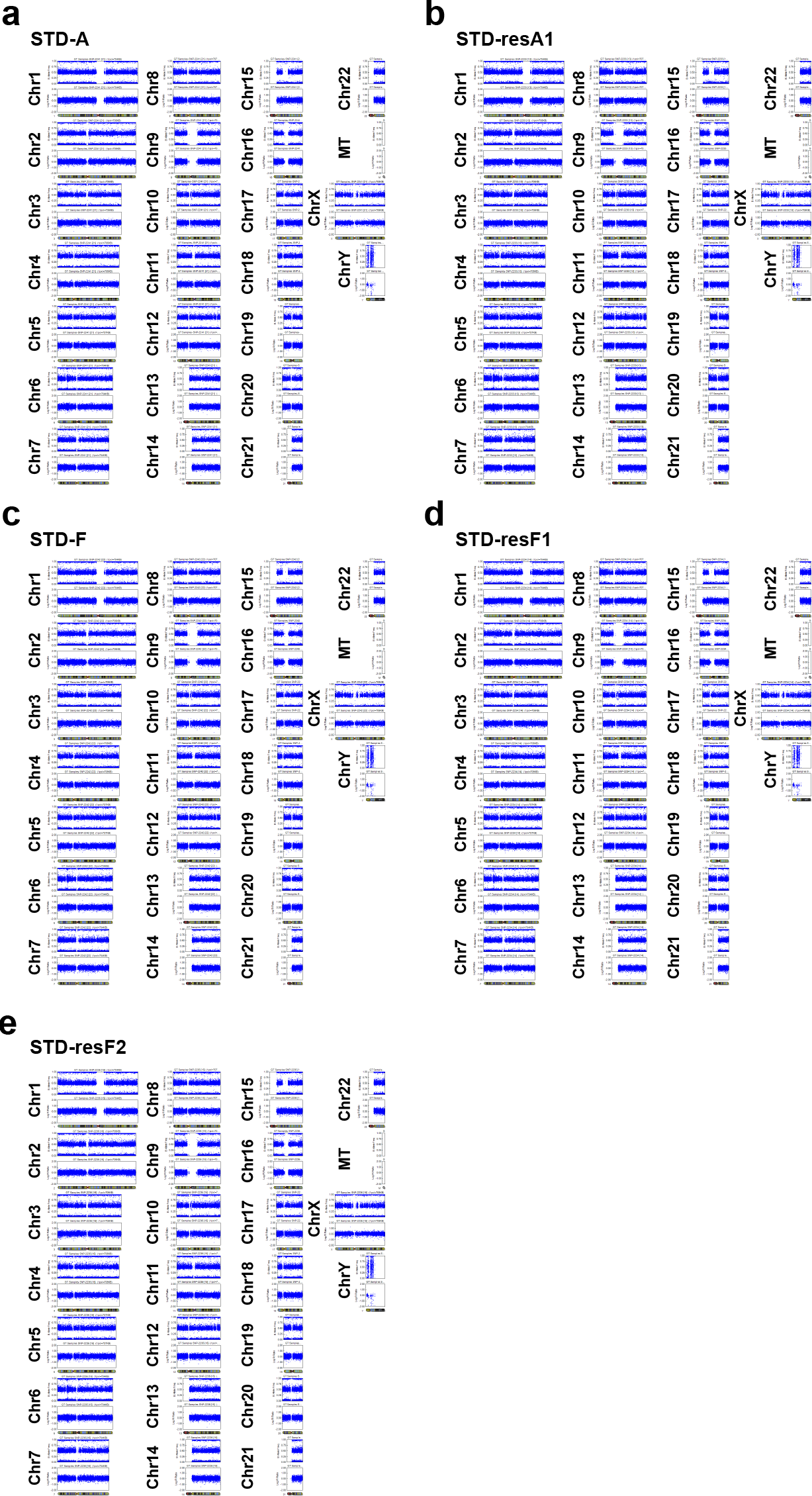
Evaluation of patient and rescued iPSCs. **a**, **b**, Resulting karyograms from SNP array analysis of STD patient iPSC clone A (STD-A) and corresponding rescued iPSC line (STD-resA1). **c-e**, Karyograms from SNP array analysis of STD patient iPSC clone F (STD-F) and corresponding rescued iPSC lines (STD-resF1/F2). No de novo CNVs were detected following gene editing and subcloning. These figures were created by Illumina Genome Viewer (IGV) (Version 1.9.0) on Illumina GenomeStudio V2011.1 with Human:Build 37 genome.

## EXTENDED DATA FILES

**Extended Data Movie 1 | 3D-synchronization assay for wild-type PSM.**

Bright field view (left) and *HES7* luciferase reporter images (right). Representative data of three independent experiments are shown. Scale bar: 500 μm.

**Extended Data Movie 2 | Calcium imaging of contracting DM-derived muscle.** Representative movies of dermomyotome (DM) derived human skeletal muscle. GCaMP reporter line activity (green fluorescence) indicating calcium influx into contracting muscle cells. Magnified view of contracting muscle cells showing concomitant calcium activity (right side of movie panel). Scale bar: 100 μm.

**Extended Data Movie 3 | 3D-synchronization assay for knock out PSMs.**

HES7-reporter activity is shown for WT and *DLL3*-, *LFNG*-, *MESP2*-knock-out PSMs. Representative data of three independent experiments are shown. Scale bar: 500 μm.

**Extended Data Table 1 | RNA-seq analysis of human PSM and derivatives.** Expressed genes arranged into six major expression clusters/groups corresponding to the six distinct differentiation and developmental stages analyzed.

**Extended Data Table 2 | List of oscillating human segmentation clock genes.** Complete list of all phase and anti-phase oscillating genes (high stringency cut-off) identified by ARSER algorithm (for details see Methods section).

**Extended Data Table 3 | Pathway-analysis of identified oscillating genes.**

Complete results of pathway and GO analyses for phase and anti-phase oscillating genes.

**Extended Data Table 4 | Recombinant proteins & small molecules used in this study.** List of utilized recombinant human proteins (4.1) and small molecule agonists and inhibitors (4.2).

**Extended Data Table 5 | Primers used in this study.**

List of utilized qRT-PCR primers for differentiation and oscillation assays (5.1), qRT-PCR primers for iPSC quality control (5.2), exon-specific primers for genotyping (5.3), oligos for sgRNA cloning (5.4), InFusion primers for MhAX targeting vectors (5.5), PCR genotyping for MhAX targeting and excision (5.6).

**Extended Data Table 6 | Antibodies used in this study.**

List of utilized primary antibodies (6.1) and secondary antibodies (6.2) for immunostaining, and antibodies used for flow cytometric analysis (6.3).

### Extended Data Table 4 Utilized recombinant proteins and small molecules

**Extended Data Table 4.1.**
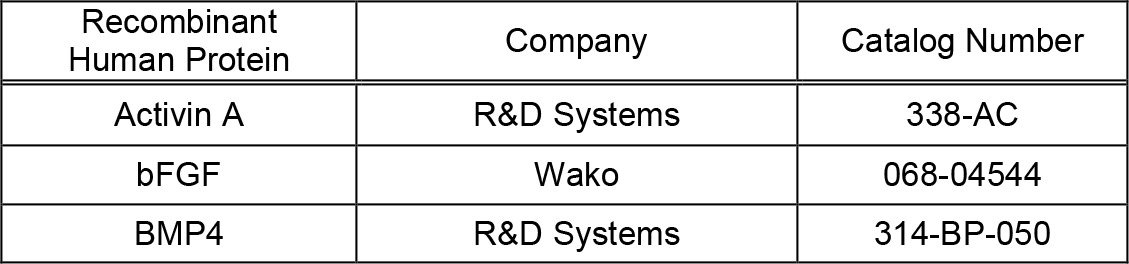
Human recombinant proteins.

**Extended Data Table 4.2.**
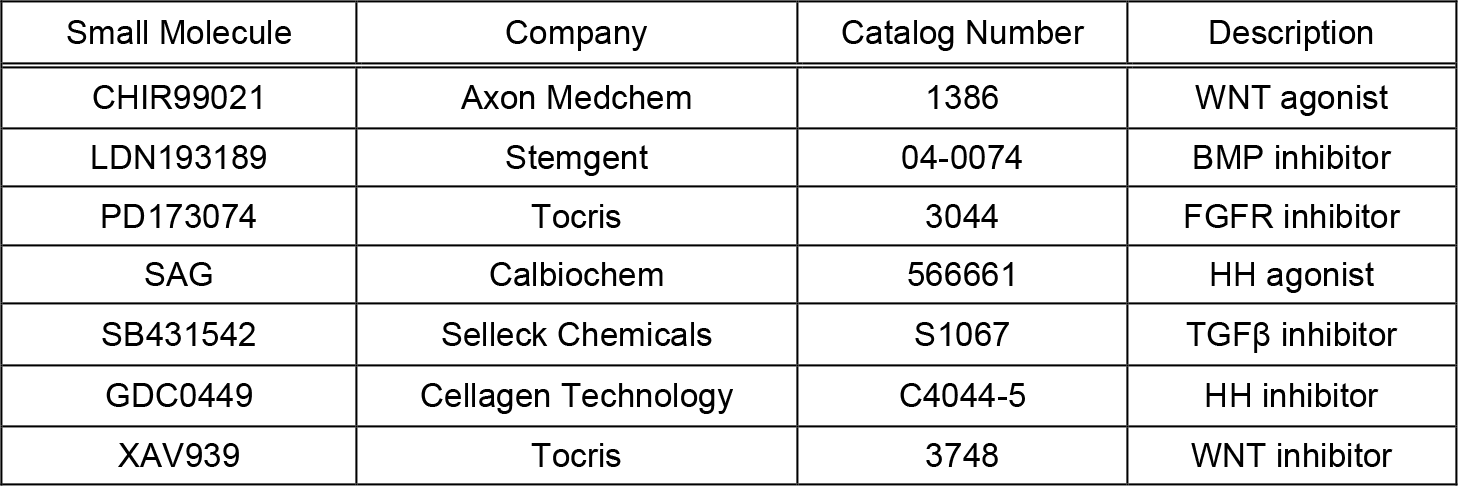
Small molecule agonists/inhibitors.

### Extended Data Table 5 Primers used in this study

**Extended Data Table 5.1.**
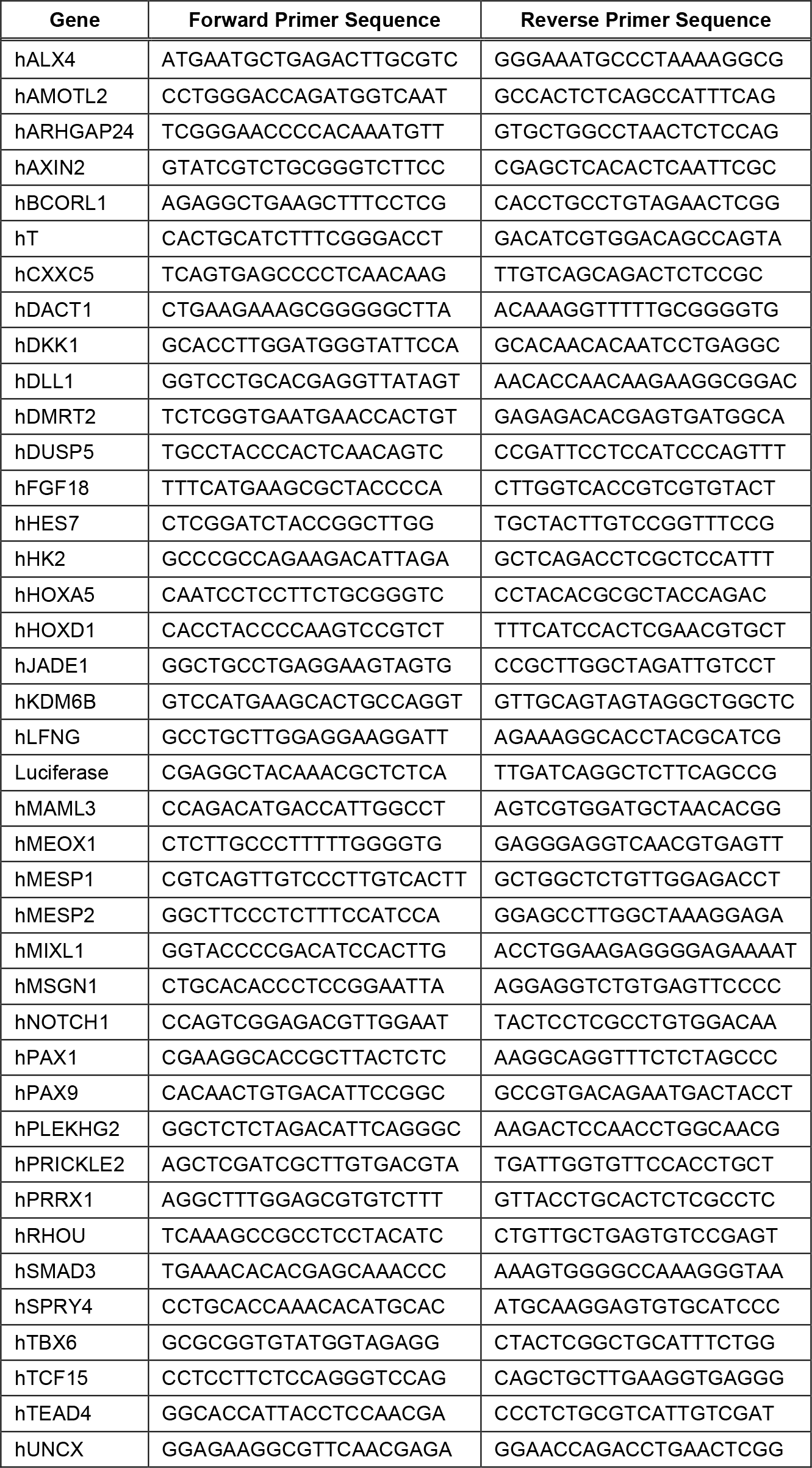
qRT-PCR primers (differentiation and oscillation)

**Extended Data Table 5.2.**
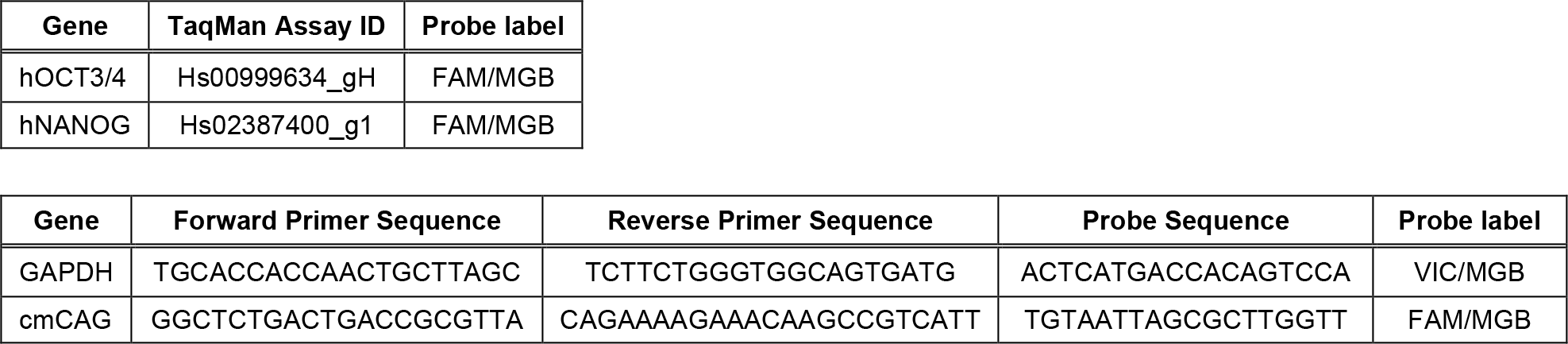
qRT-PCR primers (iPSC quality control)

**Extended Data Table 5.3.**
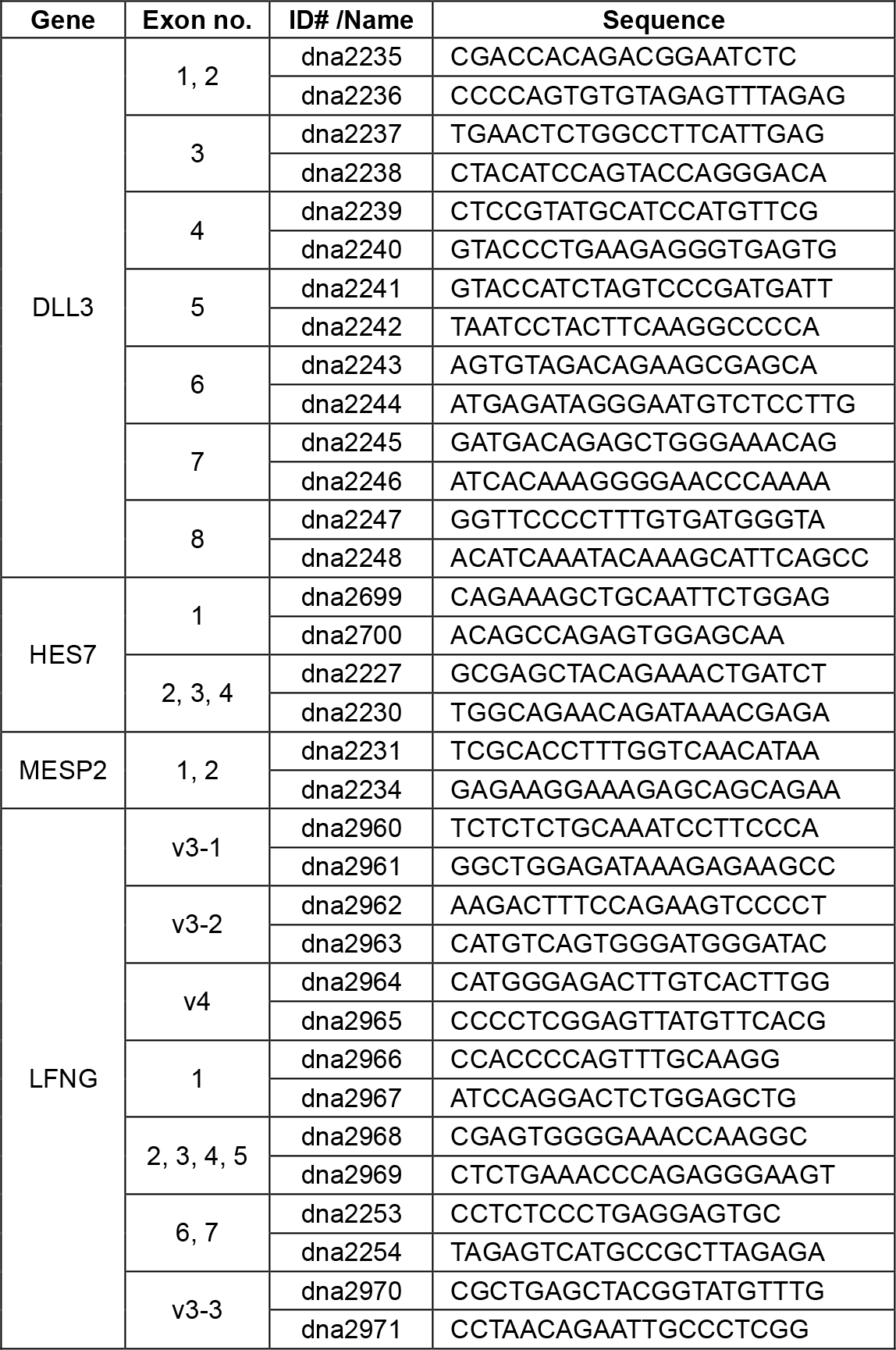
Exon genotyping primers.

**Extended Data Table 5.4.**
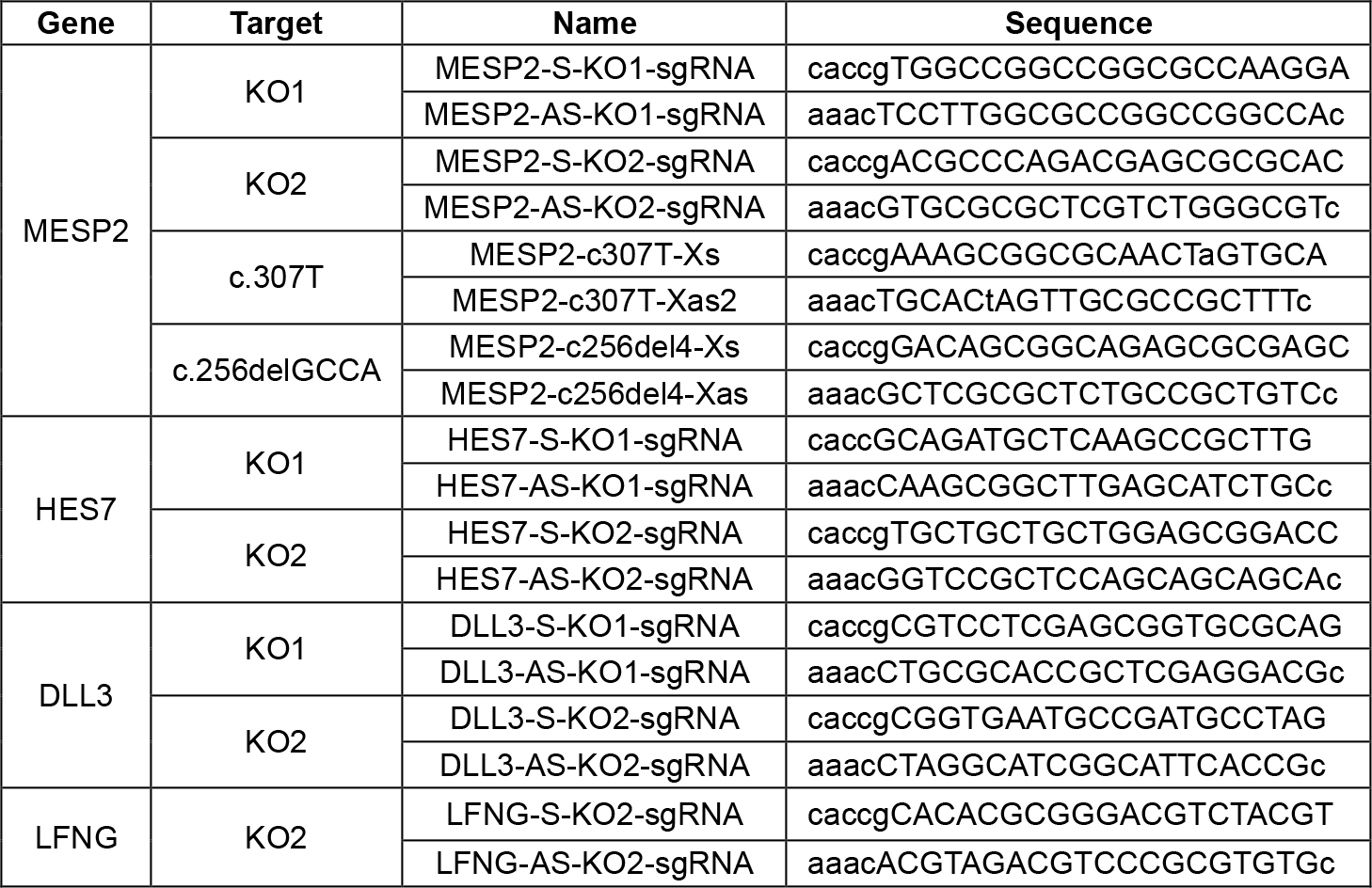
Oligos for sgRNA cloning.

**Extended Data Table 5.5.**
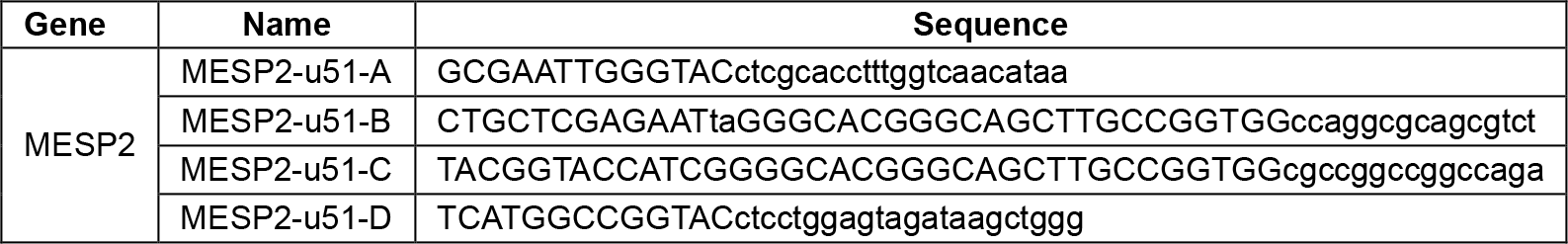
MhAX InFusion primers.

**Extended Data Table 5.6.**
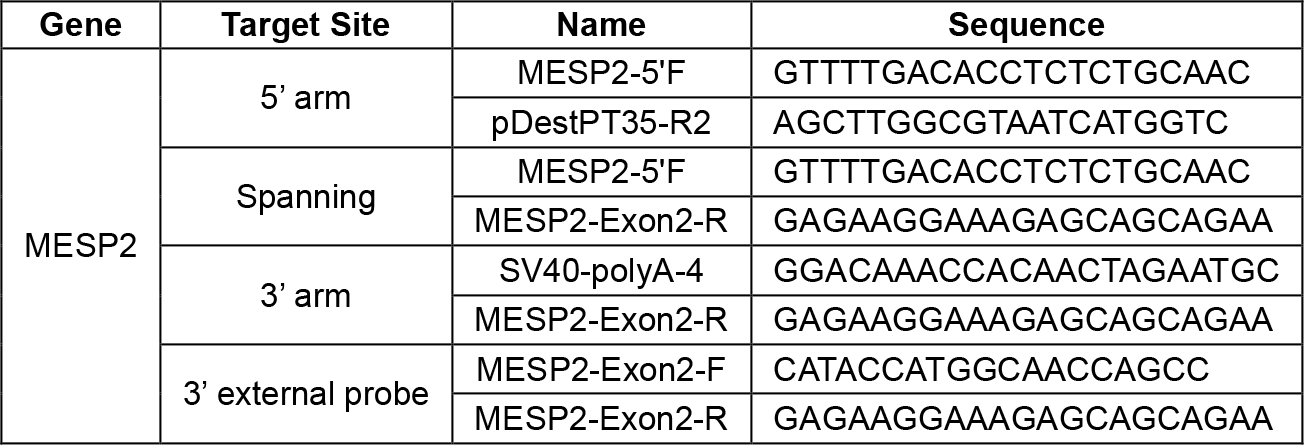
Genotyping primers.

### Extended Data Table 6 Antibodies used in this study

**Extended Data Table 6.1.**
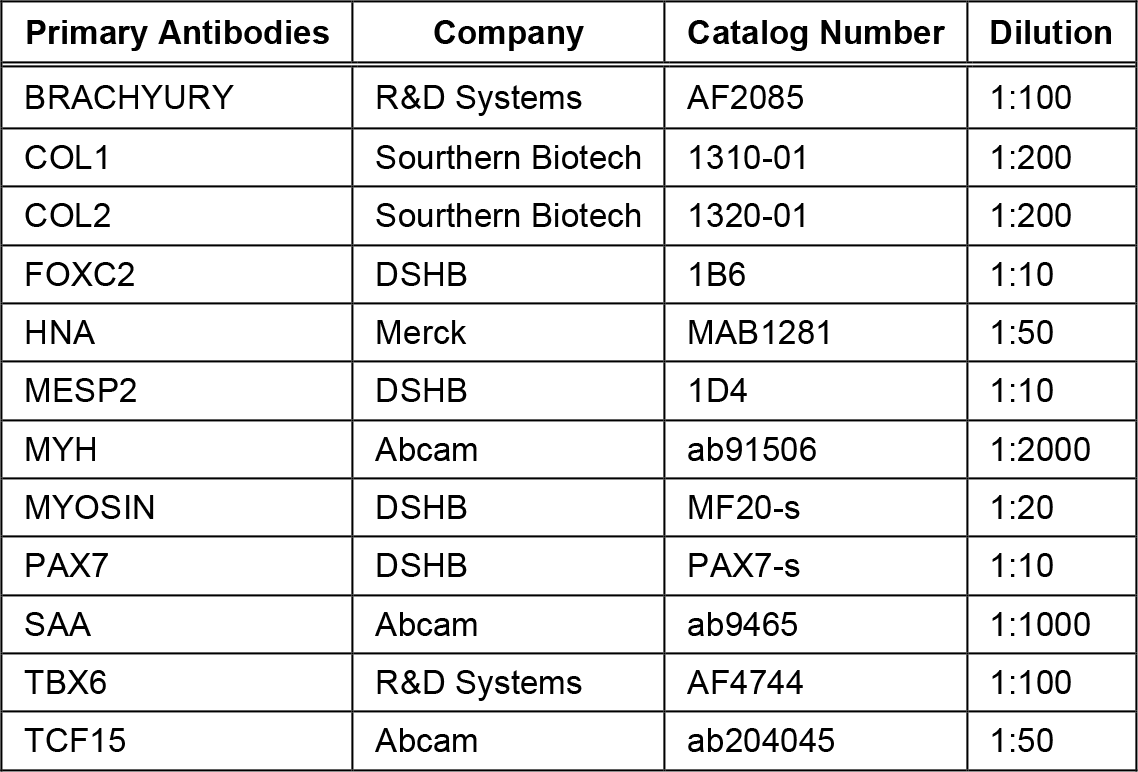
Primary antibodies used for immunostaining.

**Extended Data Table 6.2.**
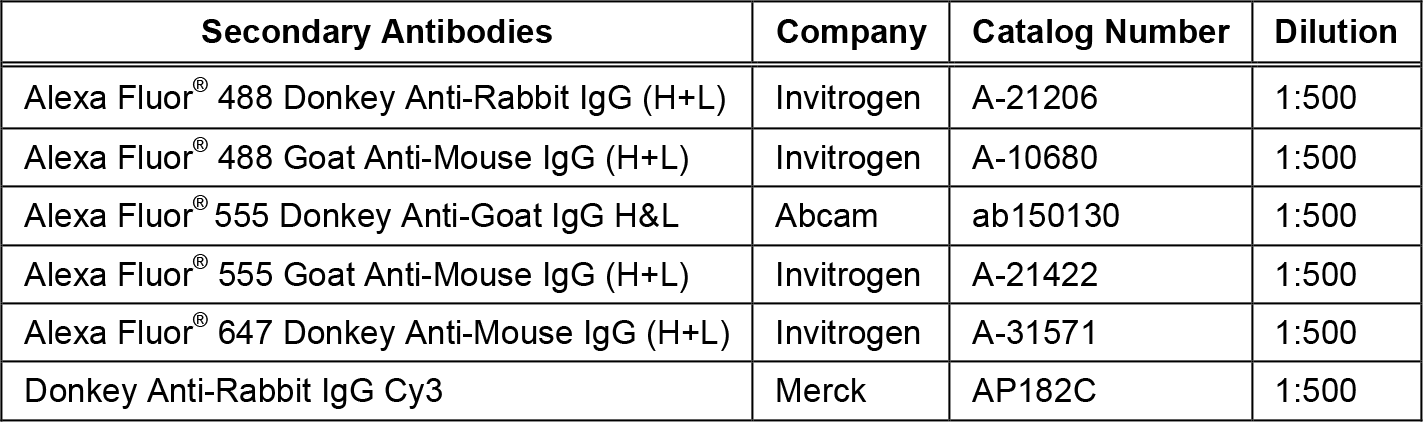
Secondary antibodies used for immunostaining.

**Extended Data Table 6.3.**
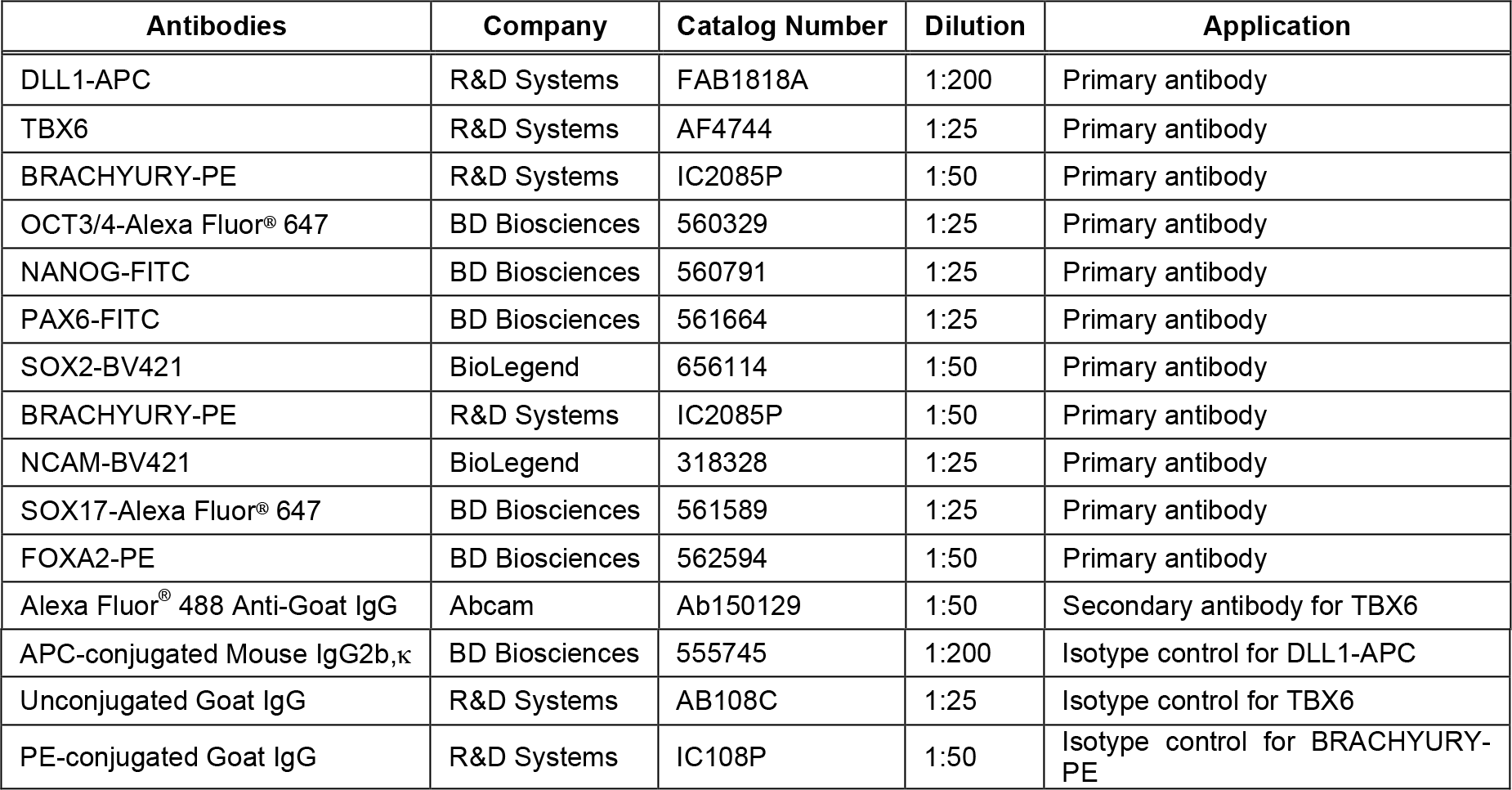
Antibodies used for flow cytometric analysis.

