## Supplemental Table 4 for "Modeling the Human Segmentation Clock with Pluripotent Stem Cells"

### Extended Data Table 4 | Utilized recombinant proteins and small molecules

### Extended Data Table 4.1 | Human recombinant proteins

| Recombinant Human Protein | Company | Catalog Number |
|---|---|---|
| Activin A | R&D Systems | 338-AC |
| bFGF | Wako | 068-04544 |
| BMP4 | R&D Systems | 314-BP-050 |

### Extended Data Table 4.2 | Small molecule agonists/inhibitors

| Small Molecule | Company | Catalog Number | Description |
|---|---|---|---|
| CHIR99021 | Axon Medchem | 1386 | WNT agonist |
| LDN193189 | Stemgent | 04-0074 | BMP inhibitor |
| PD173074 | Tocris | 3044 | FGFR inhibitor |
| SAG | Calbiochem | 566661 | HH agonist |
| SB431542 | Selleck Chemicals | S1067 | TGF $\beta$ inhibitor |
| GDC0449 | Cellagen Technology | C4044-5 | HH inhibitor |
| XAV939 | Tocris | 3748 | WNT inhibitor |
