## Supplemental Table 5 for "Modeling the Human Segmentation Clock with Pluripotent Stem Cells"

### Extended Data Table 5 | Primers used in this study

#### Extended Data Table 5.1 | qRT-PCR primers (differentiation and oscillation)

| Gene | Forward Primer Sequence | Reverse Primer Sequence |
|---|---|---|
| hALX4 | ATGAATGCTGAGACTTGCGTC | GGGAAATGCCCTAAAAGGCG |
| hAMOTL2 | CCTGGGACCAGATGGTCAAT | GCCACTCTCAGCCATTTCAG |
| hARHGAP24 | TCGGGAACCCCACAAATGTT | GTGCTGGCCTAACTCTCCAG |
| hAXIN2 | GTATCGTCTGCGGGTCTTCC | CGAGCTCACACTCAATTCGC |
| hBCORL1 | AGAGGCTGAAGCTTTCCTCG | CACCTGCCTGTAGAACTCGG |
| hT | CACTGCATCTTTCGGGACCT | GACATCGTGGACAGCCAGTA |
| hCXXC5 | TCAGTGAGCCCCTCAACAAG | TTGTCAGCAGACTCTCCGC |
| hDACT1 | CTGAAGAAAGCGGGGGCTTA | ACAAAGGTTTTTGCGGGGTG |
| hDKK1 | GCACCTTGGATGGGTATTCCA | GCACAACACAATCCTGAGGC |
| hDLL1 | GGTCCTGCACGAGGTTATAGT | AACACCAACAAGAAGGCGGAC |
| hDMRT2 | TCTCGGTGAATGAACCACTGT | GAGAGACACGAGTGATGGCA |
| hDUSP5 | TGCCTACCCACTCAACAGTC | CCGATTCCTCCATCCCAGTTT |
| hFGF18 | TTTCATGAAGCGCTACCCCA | CTTGGTCACCGTCGTGTACT |
| hHES7 | CTCGGATCTACCGGCTTGG | TGCTACTTGTCCGGTTTCCG |
| hHK2 | GCCCGCCAGAAGACATTAGA | GCTCAGACCTCGCTCCATTT |
| hHOXA5 | CAATCCTCCTTCTGCGGGTC | CCTACACGCGCTACCAGAC |
| hHOXD1 | CACCTACCCCAAGTCCGTCT | TTTCATCCACTCGAACGTGCT |
| hJADE1 | GGCTGCCTGAGGAAGTAGTG | CCGCTTGGCTAGATTGTCCT |
| hKDM6B | GTCCATGAAGCACTGCCAGGT | GTTGCAGTAGTAGGCTGGCTC |
| hLFNG | GCCTGCTTGGAGGAAGGATT | AGAAAGGCACCTACGCATCG |
| Luciferase | CGAGGCTACAAACGCTCTCA | TTGATCAGGCTCTTCAGCCG |
| hMAML3 | CCAGACATGACCATTGGCCT | AGTCGTGGATGCTAACACGG |
| hMEOX1 | CTCTTGCCCTTTTTGGGGTG | GAGGGAGGTCAACGTGAGTT |
| hMESP1 | CGTCAGTTGTCCCTTGTCACTT | GCTGGCTCTGTTGGAGACCT |
| hMESP2 | GGCTTCCCTCTTTCCATCCA | GGAGCCTTGGCTAAAGGAGA |
| hMIXL1 | GGTACCCCGACATCCACTTG | ACCTGGAAGAGGGGAGAAAAT |
| hMSGN1 | CTGCACACCCTCCGGAATTA | AGGAGGTCTGTGAGTTCCCC |
| hNOTCH1 | CCAGTCGGAGACGTTGGAAT | TACTCCTCGCCTGTGGACAA |
| hPAX1 | CGAAGGCACCGCTTACTCTC | AAGGCAGGTTTCTCTAGCCC |
| hPAX9 | CACAACTGTGACATTCCGGC | GCCGTGACAGAATGACTACCT |
| hPLEKHG2 | GGCTCTCTAGACATTCAGGGC | AAGACTCCAACCTGGCAACG |
| hPRICKLE2 | AGCTCGATCGCTTGTGACGTA | TGATTGGTGTTCCACCTGCT |
| hPRRX1 | AGGCTTTGGAGCGTGTCTTT | GTTACCTGCACTCTCGCCTC |
| hRHOU | TCAAAGCCGCCTCCTACATC | CTGTTGCTGAGTGTCCGAGT |
| hSMAD3 | TGAAACACACGAGCAAACCC | AAAGTGGGGCCAAAGGGTAA |
| hSPRY4 | CCTGCACCAAACACATGCAC | ATGCAAGGAGTGTGCATCCC |
| hTBX6 | GCGCGGTGTATGGTAGAGG | CTACTCGGCTGCATTTCTGG |
| hTCF15 | CCTCCTTCTCCAGGGTCCAG | CAGCTGCTTGAAGGTGAGGG |
| hTEAD4 | GGCACCATTACCTCCAACGA | CCCTCTGCGTCATTGTCGAT |
| hUNCX | GGAGAAGGCGTTCAACGAGA | GGAACCAGACCTGAACTCGG |

#### Extended Data Table 5.2 | qRT-PCR primers (iPSC quality control)

| Gene | TaqMan Assay ID | Probe label |
|---|---|---|
| hOCT3/4 | Hs00999634_gH | FAM/MGB |
| hNANOG | Hs02387400_g1 | FAM/MGB |

| Gene | Forward Primer Sequence | Reverse Primer Sequence | Probe Sequence | Probe label |
|---|---|---|---|---|
| GAPDH | TGCACCACCAACTGCTTAGC | TCTTCTGGGTGGCAGTGATG | ACTCATGACCACAGTCCA | VIC/MGB |
| cmCAG | GGCTCTGACTGACCGCGTTA | CAGAAAAGAAACAAGCCGTCATT | TGTAATTAGCGCTTGGTT | FAM/MGB |

#### Extended Data Table 5.3 | Exon genotyping primers

| Gene | Exon no. | ID# /Name | Sequence |
|---|---|---|---|
| DLL3 | 1, 2 | dna2235 | CGACCACAGACGGAATCTC |
| | | dna2236 | CCCCAGTGTGTAGAGTTTAGAG |
| | 3 | dna2237 | TGAACTCTGGCCTTCATTGAG |
| | | dna2238 | CTACATCCAGTACCAGGGACA |
| | 4 | dna2239 | CTCCGTATGCATCCATGTTCG |
| | | dna2240 | GTACCCTGAAGAGGGTGAGTG |
| | 5 | dna2241 | GTACCATCTAGTCCCGATGATT |
| | | dna2242 | TAATCCTACTTCAAGGCCCCA |
| | 6 | dna2243 | AGTGTAGACAGAAGCGAGCA |
| | | dna2244 | ATGAGATAGGGAATGTCTCCTTG |
| | 7 | dna2245 | GATGACAGAGCTGGGAAACAG |
| | | dna2246 | ATCACAAAGGGGAACCCAAAA |
| | 8 | dna2247 | GGTTCCCCTTTGTGATGGGTA |
| | | dna2248 | ACATCAAATACAAAGCATTCAGCC |
| HES7 | 1 | dna2699 | CAGAAAGCTGCAATTCTGGAG |
| | | dna2700 | ACAGCCAGAGTGGAGCAA |
| | 2, 3, 4 | dna2227 | GCGAGCTACAGAAACTGATCT |
| | | dna2230 | TGGCAGAACAGATAAACGAGA |
| MESP2 | 1, 2 | dna2231 | TCGCACCTTTGGTCAACATAA |
| | | dna2234 | GAGAAGGAAAGAGCAGCAGAA |
| LFNG | v3-1 | dna2960 | TCTCTCTGCAAATCCTTCCCA |
| | | dna2961 | GGCTGGAGATAAAGAGAAGCC |
| | v3-2 | dna2962 | AAGACTTTCCAGAAGTCCCCT |
| | | dna2963 | CATGTCAGTGGGATGGGATAC |
| | v4 | dna2964 | CATGGGAGACTTGTCACTTGG |
| | | dna2965 | CCCCTCGGAGTTATGTTCACG |
| | 1 | dna2966 | CCACCCCAGTTTGCAAGG |
| | | dna2967 | ATCCAGGACTCTGGAGCTG |
| | 2, 3, 4, 5 | dna2968 | CGAGTGGGGAAACCAAGGC |
| | | dna2969 | CTCTGAAACCCAGAGGGAAGT |
| | 6, 7 | dna2253 | CCTCTCCCTGAGGAGTGC |
| | | dna2254 | TAGAGTCATGCCGCTTAGAGA |
| | v3-3 | dna2970 | CGCTGAGCTACGGTATGTTTG |
| | | dna2971 | CCTAACAGAATTGCCCTCGG |

#### Extended Data Table 5.4 | Oligos for sgRNA cloning

| Gene | Target | Name | Sequence |
|---|---|---|---|
| MESP2 | KO1 | MESP2-S-KO1-sgRNA | caccgTGGCCGGCCGGCGCCAAGGA |
| | | MESP2-AS-KO1-sgRNA | aaacTCCTTGGCGCCGGCCGGCCAc |
| | KO2 | MESP2-S-KO2-sgRNA | caccgACGCCCAGACGAGCGCGCAC |
| | | MESP2-AS-KO2-sgRNA | aaacGTGCGCGCTCGTCTGGGCGTc |
| | c.307T | MESP2-c307T-Xs | caccgAAAGCGGCGCAACTaGTGCA |
| | | MESP2-c307T-Xas2 | aaacTGCACtAGTTGCGCCGCTTTc |
| | c.256delGCCA | MESP2-c256del4-Xs | caccgGACAGCGGCAGAGCGCGAGC |
| | | MESP2-c256del4-Xas | aaacGCTCGCGCTCTGCCGCTGTCc |
| HES7 | KO1 | HES7-S-KO1-sgRNA | caccGCAGATGCTCAAGCCGCTTG |
| | | HES7-AS-KO1-sgRNA | aaacCAAGCGGCTTGAGCATCTGC |
| | KO2 | HES7-S-KO2-sgRNA | caccgTGCTGCTGCTGGAGCGGACC |
| | | HES7-AS-KO2-sgRNA | aaacGGTCCGCTCCAGCAGCAGCAc |
| DLL3 | KO1 | DLL3-S-KO1-sgRNA | caccgCGTCCTCGAGCGGTGCGCAG |
| | | DLL3-AS-KO1-sgRNA | aaacCTGCGCACCGCTCGAGGACGc |
| | KO2 | DLL3-S-KO2-sgRNA | caccgCGGTGAATGCCGATGCCTAG |
| | | DLL3-AS-KO2-sgRNA | aaacCTAGGCATCGGCATTCACCGc |
| LFNG | KO2 | LFNG-S-KO2-sgRNA | caccgCACACGCGGGACGTCTACGT |
| | | LFNG-AS-KO2-sgRNA | aaacACGTAGACGTCCCGCGTGTGc |

#### Extended Data Table 5.5 | MhAX InFusion primers

| Gene | Name | Sequence |
|---|---|---|
| MESP2 | MESP2-u51-A | GCGAATTGGGTACctcgcacctttggtcaacataa |
| | MESP2-u51-B | CTGCTCGAGAATtaGGGCACGGGCAGCTTGCCGGTGGccaggcgcagcgtct |
| | MESP2-u51-C | TACGGTACCATCGGGGCACGGGCAGCTTGCCGGTGGcgccggccggccaga |
| | MESP2-u51-D | TCATGGCCGGTACctcctggagtagataagctggg |

#### Extended Data Table 5.6 | Genotyping primers

| Gene | Target Site | Name | Sequence |
|---|---|---|---|
| MESP2 | 5’ arm | MESP2-5'F | GTTTTGACACCTCTCTGCAAC |
| | | pDestPT35-R2 | AGCTTGGCGTAATCATGGTC |
| | Spanning | MESP2-5'F | GTTTTGACACCTCTCTGCAAC |
| | | MESP2-Exon2-R | GAGAAGGAAAGAGCAGCAGAA |
| | 3’ arm | SV40-polyA-4 | GGACAAACCACAACTAGAATGC |
| | | MESP2-Exon2-R | GAGAAGGAAAGAGCAGCAGAA |
| | 3’ external probe | MESP2-Exon2-F | CATACCATGGCAACCAGCC |
| | | MESP2-Exon2-R | GAGAAGGAAAGAGCAGCAGAA |
