## Supplemental Table 6 for "Modeling the Human Segmentation Clock with Pluripotent Stem Cells"

### Extended Data Table 6 | Antibodies used in this study

#### Extended Data Table 6.1 | Primary antibodies used for immunostaining

| Primary Antibodies | Company | Catalog Number | Dilution |
|---|---|---|---|
| BRACHYURY | R&D Systems | AF2085 | 1:100 |
| COL1 | Sourthern Biotech | 1310-01 | 1:200 |
| COL2 | Sourthern Biotech | 1320-01 | 1:200 |
| FOXC2 | DSHB | 1B6 | 1:10 |
| HNA | Merck | MAB1281 | 1:50 |
| MESP2 | DSHB | 1D4 | 1:10 |
| MYH | Abcam | ab91506 | 1:2000 |
| MYOSIN | DSHB | MF20-s | 1:20 |
| PAX7 | DSHB | PAX7-s | 1:10 |
| SAA | Abcam | ab9465 | 1:1000 |
| TBX6 | R&D Systems | AF4744 | 1:100 |
| TCF15 | Abcam | ab204045 | 1:50 |

#### Extended Data Table 6.2 | Secondary antibodies used for immunostaining

| Secondary Antibodies | Company | Catalog Number | Dilution |
|---|---|---|---|
| Alexa Fluor® 488 Donkey Anti-Rabbit IgG (H+L) | Invitrogen | A-21206 | 1:500 |
| Alexa Fluor® 488 Goat Anti-Mouse IgG (H+L) | Invitrogen | A-10680 | 1:500 |
| Alexa Fluor® 555 Donkey Anti-Goat IgG H&L | Abcam | ab150130 | 1:500 |
| Alexa Fluor® 555 Goat Anti-Mouse IgG (H+L) | Invitrogen | A-21422 | 1:500 |
| Alexa Fluor® 647 Donkey Anti-Mouse IgG (H+L) | Invitrogen | A-31571 | 1:500 |
| Donkey Anti-Rabbit IgG Cy3 | Merck | AP182C | 1:500 |

#### Extended Data Table 6.3 | Antibodies used for flow cytometric analysis

| Antibodies | Company | Catalog Number | Dilution | Application |
|---|---|---|---|---|
| DLL1-APC | R&D Systems | FAB1818A | 1:200 | Primary antibody |
| TBX6 | R&D Systems | AF4744 | 1:25 | Primary antibody |
| BRACHYURY-PE | R&D Systems | IC2085P | 1:50 | Primary antibody |
| OCT3/4- Alexa Fluor® 647 | BD Biosciences | 560329 | 1:25 | Primary antibody |
| NANOG-FITC | BD Biosciences | 560791 | 1:25 | Primary antibody |
| PAX6-FITC | BD Biosciences | 561664 | 1:25 | Primary antibody |
| SOX2-BV421 | BioLegend | 656114 | 1:50 | Primary antibody |
| BRACHYURY-PE | R&D Systems | IC2085P | 1:50 | Primary antibody |
| NCAM-BV421 | BioLegend | 318328 | 1:25 | Primary antibody |
| SOX17-Alexa Fluor® 647 | BD Biosciences | 561589 | 1:25 | Primary antibody |
| FOXA2-PE | BD Biosciences | 562594 | 1:50 | Primary antibody |
| Alexa Fluor® 488 Anti-Goat IgG | Abcam | Ab150129 | 1:50 | Secondary antibody for TBX6 |

| | | | | |
|---|---|---|---|---|
| APC-conjugated Mouse IgG2b,κ | BD Biosciences | 555745 | 1:200 | Isotype control for DLL1-APC |
| Unconjugated Goat IgG | R&D Systems | AB108C | 1:25 | Isotype control for TBX6 |
| PE-conjugated Goat IgG | R&D Systems | IC108P | 1:50 | Isotype control for BRACHYURY-PE |
